# Human Plasma-Like Medium Promotes Maturation of Human Pluripotent Stem Cell-Derived Cardiomyocytes

**DOI:** 10.1101/2025.04.24.650456

**Authors:** Xiaotian Zhang, Aaron D. Simmons, Kimberly S. Huggler, Austin K. Feeney, Jason R. Cantor, Melissa C. Skala, Sean P. Palecek

## Abstract

Maturing human pluripotent stem cell-derived cardiomyocytes (hPSC-CMs) *in vitro* is critical for advancing drug discovery and cardiotoxicity screening applications of these cells. However, the metabolic compositions of basal media used for hPSC-CM culture typically offer limited relevance to human cardiac physiology. Here, we examined how culture in Human Plasma-Like Medium (HPLM) versus conventional basal media affects the behavior of hPSC-CMs. Starting with Day 16 hPSC-CMs, we cultured cells for two weeks in either HPLM or RPMI-based media and then assessed maturation outcomes at Day 30. Compared to RPMI/B27 media containing either RPMI-defined (11.1 mM) or physiologic glucose levels (5 mM), HPLM/B27 markedly enhanced hPSC-CM maturity as evinced by concerted transcriptomic, structural, functional, and metabolic phenotypes. These effects included a higher extent of myosin heavy chain isoform switching (α-MHC to β-MHC), accelerated ventricular-specific myosin light chain isoform switching (MLC2a to MLC2v), elongated sarcomeres, increased multinucleation, enhanced calcium transient kinetics, and coordinated activation of oxidative and glycolytic metabolism. Collectively, these findings demonstrate that medium composition has substantial effects on hPSC-CM biology and also establish HPLM as a tool for driving hPSC-CM maturation *in vitro*.

**Translational Impact Statement:** HPLM was designed to more closely recapitulate the metabolic composition of human plasma and thus provides a physiological platform to promote hPSC-CM maturation. By enhancing structural, functional, and metabolic maturity, HPLM-cultured hPSC-CMs better approximate cardiac physiology, positioning them as improved models for cardiovascular disease research, drug-induced cardiotoxicity screening, and personalized therapeutic testing. This medium can integrate with existing maturation strategies, accelerating the translation of basic cardiac research into clinically predictive tools for the drug development pipeline.

**Graphical Abstract:** 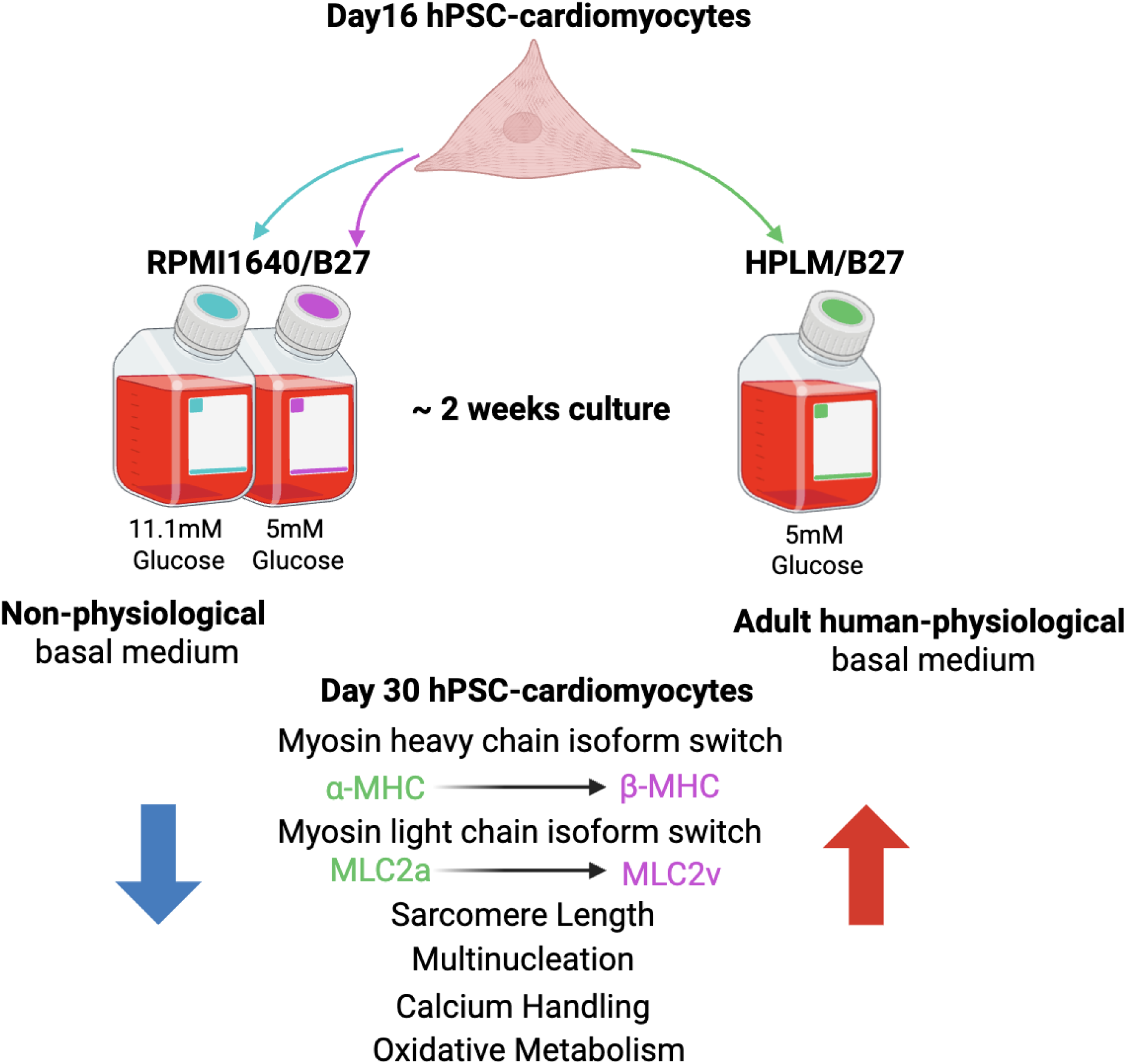

## 1. Introduction

Human pluripotent stem cell-derived cardiomyocytes (hPSC-CMs) are advancing cardiovascular research by enabling patient-specific insights into genetic disorders, drug development, and regenerative therapies^1–3^. A critical priority lies in improving preclinical drug screening to address systemic inefficiencies in cardiovascular drug discovery^4,5^. Current approaches relying on animal models poorly recapitulate aspects of human cardiac physiology^6^. While hPSC-CMs offer human-relevant alternatives, their utility is constrained by immature phenotypes that inadequately recapitulate adult cardiomyocyte function^1,2,7^, which could lead to misclassification of compounds with cardiotoxic effects as safe^4,8^. Refining hPSC-CM maturation is thus essential to bridge the translational gap between preclinical models and clinical outcomes.

Cardiomyocyte maturation requires coordinated structural, functional, and metabolic remodeling. Structurally, adult cardiomyocyte maturation involves sarcomere elongation ^9,10^, increased multinucleation ^7,11^, and sequential isoform switching of key sarcomere genes^2,7,12^. These include transitions of the early developmental to matured isoforms of myosin heavy chain as early as week 5 of gestation (*MYH6* to *MYH7*) ^1,13^, myosin light chain (*MYL7* to *MYL2*) later in ventricular cardiomyocyte development ^1,14^, and troponin I (*TNNI1* to *TNNI3*) during the pre-natal to post-natal transition ^15,16^. These coordinated isoform transitions are critical for sarcomere stabilization and contractile efficiency, which serve as essential structural and functional benchmarks to evaluate maturation in hPSC-CM models^7^. Functionally, maturation involves robust calcium handling, characterized by rapid, high-amplitude Ca²⁺ transients mediated by coordinated sarcoplasmic reticulum (SR) coupling and increased gene expression of calcium-handling machinery^7,13,14^.

Metabolically, maturation shifts energy production from glycolysis to an oxidative metabolic state, marked by increased mitochondrial volume^2,17,18^. To address these maturation deficits, strategies such as electrical stimulation^19^, 3D engineered tissues^20^, mechanical loading^21^, and co-culture with non-myocytes^22^ have been employed to enhance maturation. However, these approaches face scalability limitations or require specialized equipment. In contrast, culture media optimization by tuning nutrient composition, growth factors, and metabolic substrates offers a scalable, cost-effective alternative to systematically drive maturation while maintaining experimental simplicity^23–27^.

The traditional hPSC-CM differentiation and culture medium, RPMI 1640/B27, was not designed to support the unique physiological needs of cardiomyocytes^24^. RPMI 1640 contains supraphysiologic glucose (11.1 mM) and 19 amino acids at concentrations that in large part markedly differ compared to those in biofluids such as plasma while also lacking a variety of other macronutrients such as lipids and ketone bodies^28^. RPMI also contains sub-physiological levels of calcium (0.42 mmol/L), which in turn may hinder excitation-contraction coupling and myofibrillogenesis^29–31^. To overcome these limits, there have been media development efforts to more closely mimic the mature cardiac microenvironment. For example, a recent study reported the design of a medium based on modified DMEM to better address adult cardiomyocyte requirements, including the incorporation more relevant levels of glucose (3 mM) and calcium (1.8 mM), inclusion of various fatty acids, and key adult cardiac supplements (L-carnitine, taurine, and creatine)^24^. While this hybrid formulation led to improved maturation, it was still based on a traditional medium that does not recapitulate adult plasma metabolite concentrations.

The importance of an adult environment is supported by transplantation studies showing that hPSC-CMs achieve greater maturation in adult murine hosts than in neonatal settings ^32^. To better mirror the adult biochemical milieu and test its potential for maturing hPSC-CM *in vitro*, we tested Human Plasma-Like Medium (HPLM) ^33^, a synthetic physiologic medium systematic formulation that contains over sixty metabolites and small ions at concentrations reflective of those in adult human plasma.

To evaluate the maturation-enhancing impact of culture in HPLM, we compared HPLM/B27 to RPMI/B27 containing either RPMI-defined (11.1 mM) or HPLM-defined (5 mM) glucose levels over two-week cultures of hPSC-CMs starting at a fully-differentiated but immature phenotype achieved by Day 16 in the standard GiWi protocol ^34,35^. Using transcriptomic, structural, functional, and metabolic analyses, we demonstrated that HPLM promotes enhanced maturation linked to differences in metabolite availability that extend beyond glucose. By integrating these multidimensional assessments, this study establishes HPLM as a physiologically relevant platform for advancing hPSC-CM maturity, offering a resource for improving models of cardiac development, disease, and therapeutic testing.

## 2. Materials and methods

### 2.1 hPSC-CPC and hPSC-CM differentiation

Differentiation of WTC11, H9, and IMR90-4 hPSCs to cardiac progenitor cells (CPC) was performed following a previous protocol ^36^. Briefly, hPSCs were dissociated with Accutase and resuspended in mTeSR1 containing 5 μM Y-27632. Cells were then plated onto Matrigel-coated 12-well plates and incubated at 37°C with 5% CO₂. After 24hr, the medium was replaced with fresh mTeSR1. Differentiation was initiated on Day 0 in RPMI1640 with 2% B27 minus insulin (R/B-) and 6-12 μM CHIR99021. At Day 2, the medium was changed to R/B-containing 5 μM IWP2. By Day 4, cultures were transferred to R/B-alone. On Day 5/6, CPCs were dissociated with Accutase, pooled, and cryopreserved as described in a previous protocol ^36^. Cryopreserved hPSC-CPCs (Day 5/6) were thawed and reseeded following previous methods for differentiation to CMs^36^. Media changes with RPMI/B27 commenced on Day 5/6, continuing every 3 days until harvest at Day 16 for flow cytometry and downstream experiments.

### 2.2 Flow cytometry

Cardiomyocyte purity was quantified on Day 16 by flow cytometry for cardiac troponin T (cTnT), adapted from published protocols ^36^. Cells were dissociated using Accutase, fixed with 1% paraformaldehyde, and preserved in 90% methanol at −20°C. ∼1 million cells per sample were washed in flow buffer 1 (FB: 0.5% BSA in DPBS) and permeabilized with 0.1% Triton X-100 in FB1. Fixed cells were incubated overnight at 4°C with a 1:1000 mouse IgG1 anti-cTnT primary antibody in flow buffer 2 (0.5% BSA/DPBS + 0.1% Triton X-100), followed by three FB1 washes. Secondary staining was performed for 1 hr in the dark at room temperature using 1:1000 Alexa Fluor 488-conjugated anti-mouse IgG1 in FB2. After final washes, cells were resuspended in 300 μL FB and analyzed on a BD Accuri C6 Plus flow cytometer. Undifferentiated hPSCs underwent identical staining procedures as negative controls.

### 2.3 RNA sample collection and purification

Samples were washed once with DPBS and incubated for 1 min in 500 μL ice-cold Trizol reagent. Cells were detached, transferred to microcentrifuge tubes, flash-frozen in liquid nitrogen, and stored at −80°C. RNA extraction was performed using Trizol, followed by purification and concentration via Zymo Direct-zol MiniPrep Plus columns, including on-column DNase I treatment following manufacturer guidelines.

### 2.4 RNAseq

Extracted total RNA was validated for integrity and purity. Sequencing libraries were prepared with the Takara SMARTer Stranded Total RNA Kit – HI Mammalian and sequenced on an Illumina NovaSeq6000. Raw FASTQ files were aligned to the human genome (hg38 + decoy) with STAR v2.5.3a ^37^, and gene-level counts were quantified using featureCounts v2.0.3^38^. Differential expression analysis was conducted in R v4.1.3 with DESeq2^39^, applying variance-stabilized counts for principal component analysis (PCA) and filtering results at a false discovery rate (p-adj < 0.05). GSEA was used for gene ontology (GO) enrichment analyses. Heatmaps were created by MetaboAnalyst (version 6.0)^40^.

### 2.5 Reverse transcription, quantitative polymerase chain reaction (RT-qPCR)

cDNA was produced with the Qiagen Omniscript RT Kit, Oligo dT(20) primers, and RNaseOUT following the manufacturer’s protocol. qPCR reactions followed by SYBR Green Master Mix with target-specific primers (Table S2). Amplification was conducted on an AriaMx Real-Time PCR System. Data were normalized using the ΔCT method against the geometric mean of three reference genes (*ZNF384*, *DDB1*, *EDF1*) validated in previous work ^41^.

### 2.6 Immunostaining for cardiomyocyte structural analysis

Day 16 hPSC-CMs were cultured for two weeks in either glucose-matched RPMI/B27 (5 mM) or HPLM/B27, then replated into ibidi 96-well imaging plates at 60,000–70,000 cells/well with corresponding media for 7 days before staining. Cells were washed with DPBS, fixed in 1% paraformaldehyde, and blocked with 0.5–1% BSA/DPBS before storage at 4°C. Wells were incubated overnight at 4°C with primary antibodies in FB2 under gentle agitation, followed by secondary antibodies (1 h, RT) and Hoechst 33342 (1:5000). Imaging was performed on a microscope equipped with a Lumencor Aura III light engine.

For myosin heavy chain isoform, 1:100 rb-IgG-anti-MYH7 and 1:100 ms-IgG1-anti-MYH6 were stained for primary antibodies, with 1:1000 Alexa Fluor 647 anti-rabbit IgG, and 1:1000 Alexa Fluor 488-conjugated anti-mouse IgG1 for secondary antibodies. β-MHC(MYH7)+ and α-MHC(MYH6)+ cardiomyocytes were quantified at 20x magnification. For myosin light chain isoform, 1:200 mouse IgG2b anti-MLC2A and 1:200 rabbit IgG anti-MLC2V were stained for primary antibodies, with 1:1000 Alexa Fluor 488 anti-mouse IgG2b and Alexa Fluor 647 anti-rabbit IgG as secondary antibodies. MLC2V+ and MLC2A+ cardiomyocytes were quantified at 20x magnification. For the sarcomere, 1:1000 mouse IgG1 anti-α-actinin was stained as the primary antibody with 1:1000 Alexa Fluor 647 anti-mouse IgG1 as the secondary antibody.

Sarcomeres were analyzed at 40x magnification in SotaTool^42^.

### 2.7 Beat rate analysis

Phase-contrast video acquisition was performed at 20x magnification on a Nikon Ti2e with an ORCA-Flash4.0 camera (Hamamatsu C13440-20CU). Videos were acquired at 50 frames per second for 20 sec. MUSCLEMOTION (version 1.1)^43^ was used to quantify the beat rate.

### 2.8 Intracellular calcium measurement

Day 16 hPSC-CMs were replated onto Matrigel-coated ibidi 96-well imaging plates at 100,000–166,000 cells/well and cultured for two weeks in specified media. On Day 30, intracellular calcium dynamics were assessed using the FLIPR® Calcium 6 Assay Kit. Cells were incubated with the fluorescent Calcium 6 probe following the manufacturer’s instructions, followed by replacement with fresh media (RPMI_glcM/B27 or HPLM/B27) and stabilization in a 37°C 5% CO₂ incubator for 30 min. Cells were imaged on a Nikon Ti2e equipped with a 470 nm excitation 515/30 nm emission filter and an ORCA-Flash4.0 camera. For each well, 20-s videos were captured at 20x magnification, focusing on 20 synchronously beating cells per video. Three to four well replicates were analyzed per cell line. Fluorescence intensity traces were derived by averaging signals from 20 cells from the first calcium peak, subtracting background intensity, and normalizing to baseline (ΔF/F₀ = (F − F₀)/F₀). Calcium transient kinetics were calculated from normalized traces: Upstroke velocity: Rate of fluorescence increase during calcium release (Δ(ΔF/F₀)/Δt). Downstroke velocity: Rate of fluorescence decay during calcium reuptake (Δ(ΔF/F₀)/Δt).

### 2.9 Seahorse analysis

Mitochondrial respiration and basal glycolytic activity in hPSC-CMs were evaluated using a Seahorse XF96 extracellular flux analyzer. Cells were replated onto Matrigel-coated XF96 Cell Culture Microplates at 60,000–70,000 cells per well on Day 16 and cultured for two weeks in either glucose-matched RPMI/B27 or HPLM/B27. On day 30, cells were equilibrated for 1 hr at 37°C (non-CO₂) in Seahorse RPMI basal medium supplemented with 5 mM glucose, 1 mM sodium pyruvate, and 2 mM glutamine.

Mitochondrial stress was induced via sequential injections of 1.5 μM oligomycin (ATP synthase inhibitor), 1 μM FCCP (uncoupler), and 0.5 μM rotenone/antimycin A (Complex I/III inhibitors). Post-assay, cell numbers were normalized using Hoechst 33342 and imaged at 10x magnification. 9 independent differentiations were performed for both media conditions, with each differentiation including 5 well replicates averaged for OCR and ECAR quantification. OCR and ECAR were quantified as follows: basal OCR (average baseline OCR, time points 1–3) reflected steady-state oxidative phosphorylation; maximal OCR (FCCP-stimulated OCR minus residual post-rotenone/antimycin A OCR) represented peak respiratory capacity; basal ECAR (average baseline ECAR, time points 1–3) indicated glycolytic activity. Metabolic phenotypes were assessed using the basal OCR/ECAR ratio to evaluate oxidative phosphorylation-glycolysis balance under baseline conditions.

### 2.10 mtDNA-to-nDNA ratio analysis

qPCR was performed to determine the ratio between mitochondrial and nuclear DNA as previously described^44^. Mitochondrial gene expression was normalized for nuclear gene expression, and the ratio was normalized to the value of the Day 16 hPSC-CMs for each media treatment group.

## 2.11 Autofluorescence imaging of NAD(P)H

Cells were replated onto Matrigel-coated ibidi 96-well plates at 60,000–70,000 cells/well on Day 16 and cultured for two weeks in glucose-matched RPMI/B27 or HPLM/B27.

Before imaging, cells were incubated with 10 nM TMRE for 30 min at 37°C to label mitochondria. Fluorescence lifetime imaging (FLIM)^45^ was performed using a Bruker Ultima two-photon system equipped with an Insight DS+ tunable laser and Swabian time tagger for photon counting. Imaging parameters included 2.0 mW laser power at the sample, 512 × 512-pixel resolution (40×/1.15 NA objective, 2.5× zoom), 3.6 µs pixel dwell time, and 1.7 s integration. Both NAD(P)H and TMRE were excited at 750 nm while emission was collected through separate PMTs for NAD(P)H (400-480 nm) and TMRE (565-615 nm).

### 2.12 Statistical analysis

RNAseq data were generated from four independent wells for each media condition. Statistical tests for each experiment are detailed in the figure legends.

## 3. Results

### 3.1 HPLM promotes maturation of cardiac transcriptional programs in hPSC-CMs

To evaluate whether HPLM can enhance hPSC-CM maturation beyond RPMI1640, we employed bulk RNA sequencing (RNAseq) to assess the global transcriptomic changes across different media conditions. Cardiac progenitor cells (CPCs) were generated from WTC11 iPSCs using the GiWi protocol ^34,35^ and were harvested and cryopreserved on Day 5. This cryopreservation step was implemented to bank homogeneous CPCs with high differentiation potency, ensuring consistent differentiation into high-purity cardiomyocyte populations (>80%) at Day 16 (Figure S1). After differentiating CPCs into hPSC-CMs, on Day 16 cells were harvested for bulk RNAseq as a baseline control. The remaining cells were randomized into three media conditions: the standard differentiation protocol formulation of RPMI/B27 (11.1 mM glucose), RPMI_glcM/B27 (RPMI with HPLM-defined 5 mM glucose), and HPLM/B27, for two weeks of culture with media refreshed every three days. On Day 30, cells from all groups were harvested for RNAseq analysis (Figure 1A).

**Figure 1.**
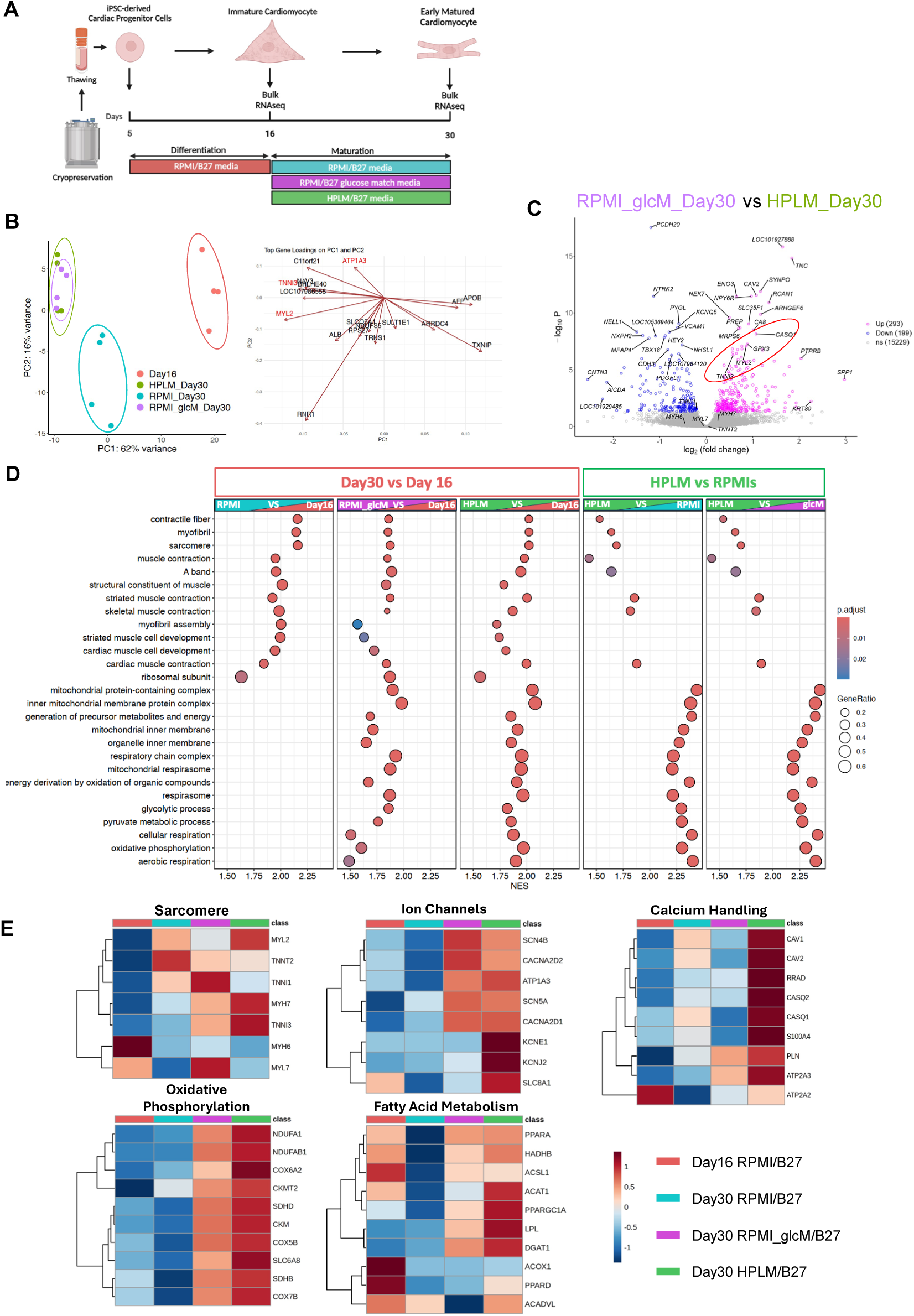
HPLM enhances the expression of cardiac physiological and metabolic genes in hPSC-CMs. (A) Schematic of the RNA-seq experimental design. Day 16 WTC11 hPSC-CMs were randomized into three media conditions: RPMI/B27, RPMI_glcM/B27 (RPMI/B27 with 5 mM glucose, concentration matched to HPLM), and HPLM/B27 and cultured for 2 weeks. RNA was extracted and sequenced from four biological replicates per condition on Day 16 and Day 30. (B) Principal component analysis (PCA) of transcriptomic profiles. Left: PCA plot of hPSC-CMs cultured under four conditions: Day 16 hPSC-CMs cultured in RPMI/B27 and Day 30 hPSC-CMs maintained in RPMI/B27, RPMI_glcM/B27, or HPLM/B27 between Day 16 and Day 30. Each dot represents a biological replicate for RNAseq, color-coded as indicated. Right: Biplot of top loading genes for PC1 and PC2. Genes associated with cardiac physiology are highlighted in red. (C) Differential expression analysis of Day 30 hPSC-CMs in RPMI_glcM/B27 versus HPLM/B27. Differentially expressed genes (adjusted P < 0.05, DESeq2 Wald test with Benjamini-Hochberg correction) are highlighted, with fold change in expression of upregulated and downregulated genes in HPLM/B27 relative to RPMI_glcM/B27. (D) Dot plots of top upregulated Gene Ontology (GO) terms associated with differentially expressed genes, summarized by gene set enrichment analysis (GSEA) across five comparisons: (1) Day 30 RPMI/B27 vs. Day 16, (2) Day 30 RPMI_glcM/B27 vs. Day 16, (3) Day 30 HPLM/B27 vs. Day 16, (4) Day 30 HPLM/B27 vs. Day 30 RPMI/B27, and (5) Day 30 HPLM/B27 vs. Day 30 RPMI_glcM/B27. Dot size reflects the number of genes associated with a pathway; color indicates adjusted p-value. NES: Normalized enrichment score. (E) Heatmap of key genes involved in aspects of cardiomyocyte maturation: sarcomere, ion channels, calcium handling, oxidative phosphorylation, and fatty acid metabolism. Media conditions (column) are color-coded from left to right: Day 16, Day 30 RPMI/B27, Day 30 RPMI_glcM/B27, and Day 30 HPLM/B27. Expression values are z-score-normalized transcripts per million (TPM).

Principal component analysis (PCA) revealed that two weeks of culture in RPMI/B27 induced a transcriptomic shift in hPSC-CMs along PC1, and this shift was further enhanced by physiologic glucose in both RPMI_glcM/B27 and HPLM/B27 media (Figure 1B). The PCA biplot highlights the contribution to PC loadings of key cardiac maturation-related genes, such as *TNNI3, MYL2* and *ATP1A3,* which were directed toward the RPMI_glcM/B27 and HPLM/B27 clusters. This suggests that prolonged culture and physiologic glucose promoted a more mature transcriptomic profile in hPSC-CMs. Although PCA did not fully distinguish Day 30 hPSC-CMs cultured in RPMI_glcM/B27 from those in HPLM/B27, differential gene expression analysis identified distinct expression profiles between these media conditions (Figure 1C and Figure S2A). Notably, *MYL2* and *TNNI3* were significantly enriched in the HPLM/B27 group (Figure 1C), which suggests that hPSC-CMs treated with HPLM/B27 exhibited the most mature cardiac-like transcriptomic profile among these conditions, given the roles of these isoforms in cardiac maturation ^46–50^.

To investigate the biological significance of these transcriptomic changes, gene set enrichment analysis (GSEA) was performed, and the top upregulated gene ontology (GO) terms across comparisons were visualized as dot plots (Figure 1D and Figure S2B).

Consistent with the PCA results, two-week culture and differential glucose levels emerged as key drivers of transcriptomic remodeling. Prolonged culture primarily upregulated GO terms associated with cardiac physiology (e.g., myofibril, sarcomere, and muscle contraction), while physiologic glucose predominantly enriched metabolic processes (e.g., mitochondrial metabolism and glycolysis). Notably, the HPLM/B27 condition amplified these effects when compared to RPMI/B27 and RPMI_glcM/B27, upregulating GO terms linked to cardiac physiology and metabolism (Figure 1D). This suggests that while physiologic glucose can activate cardiac metabolic maturation during extended culture, other HPLM components contribute to further enhancing cardiac maturation.

Complementing these pathway-level findings, differential expression analysis revealed coordinated transcriptional signatures of maturation in HPLM/B27-treated hPSC-CMs (Figure 1E). Structural maturation was marked by the upregulation of adult sarcomere isoforms (*MYH7, MYL2*, *TNNI3*) and the downregulation of earlier developmental isoforms (*MYH6*, *MYL7*, *TNNI1*). Enhanced calcium handling was supported by elevated expression of *CASQ2*, *PLN*, and ion channel genes such as *KCNJ2* and *SCN5A*, which are critical for electrophysiological maturation. Metabolic reprogramming was evidenced by upregulation of mitochondrial respiration genes (*NDUFA1*, *COX7B*, *SDHB*) and fatty acid oxidation markers (*PPARAGC1A*, *ACADVL*), consistent with a shift toward oxidative metabolism characteristic of adult cardiomyocytes. Collectively, these gene expression signatures demonstrate that while physiologic glucose RPMI/B27 partially drives metabolic remodeling, HPLM/B27 uniquely enhances a functionally integrated maturation transcriptomic program, combining structural, calcium-handling, and metabolic adaptations.

Columns (labeled H for H9, I for IMR90-4, or W for WTC11) represent independent differentiations for each cell line. Rows represent individual genes with the color scale indicating the Z-score of the qPCR results (ΔCq). (D) Principal component analysis (PCA) of the qPCR results from Figure 2C. Each dot represents an independent differentiation (labeled H for H9, I for IMR90-4, or W for WTC11) under one of the three media conditions. Points are colored by condition: pink for Day 16 differentiation baseline, purple for Day 30 RPMI_glcM/B27, and green for Day 30 HPLM/B27. (E) Top genes with medium condition-dependent expression changes, identified by Partial Least Squares Discriminant Analysis (PLS-DA) of qPCR results and ranked by Variable Importance in Projection (VIP) scores (x-axis). The color blocks on the right show the relative expression of each gene for different media treatment groups: pink for Day 16 differentiation baseline, purple for Day 30 RPMI_glcM/B27, and green for Day 30 HPLM/B27.

**Figure 2.**
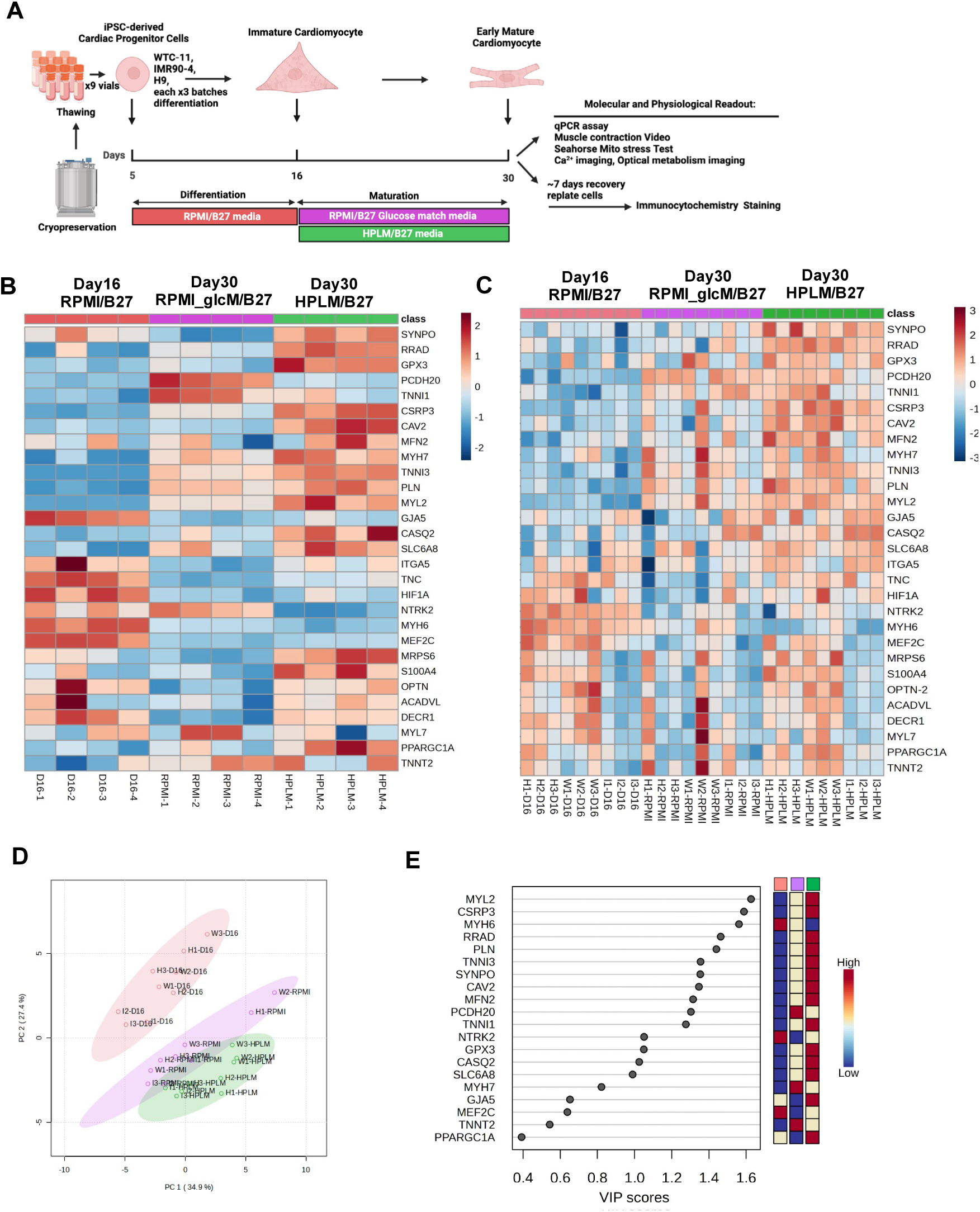
HPLM promotes maturation-associated gene expression in hPSC-CMs in multiple cell lines. (A) Schematic overview of molecular validation and functional phenotyping of hPSC-CMs across different media treatment groups. Day 16 hPSC-CMs were randomized into RPMI_glcM/B27 and HPLM/B27 media and cultured for 2 weeks before molecular and physiological assessments. qPCR analysis was performed on hPSC-CMs derived from H9, WTC11, and IMR90-4 hPSCs, with each cell line having 3 independent differentiation batches. (B) Heatmap of top differentially expressed genes identified by RNA-seq across different media treatment groups: Day 16 hPSC-CMs cultured in RPMI/B27, Day 30 hPSC-CMs cultured in RPMI_glcM /B27, and HPLM/B27. Data were generated from WTC11 hPSC-CMs, with four biological replicates (columns) per condition. Rows represent individual genes, and the color scale indicates the Z-score of TPM-based expression. (C) Heatmap of qPCR validation of differentially expressed genes identified by RNA-seq across media treatment groups: Day 16 hPSC-CMs cultured in RPMI/B27, and Day 30 hPSC-CMs cultured in RPMI_glcM/B27, and HPLM/B27.

To validate the RNA-seq findings and assess reproducibility across cell lines and differentiation batches, qPCR analysis was performed on hPSC-CMs derived from three hPSC lines (H9, IMR90-4, and WTC11), with three independent differentiations of each line (Figure 2A). We compared day 30 HPLM/B27 and RPMI_glcM/B27 to a day 16 control to identify maturation effects of HPLM beyond physiologic glucose concentration. The transcriptional effects of HPLM identified by RNA-seq were generally recapitulated by qPCR in CMs from different cell lines (Figures 2B-C). Adult sarcomere isoforms *MYL2*, *MYH7*, *TNNI3* and cardiac LIM protein *CSRP3*, calcium-related *PLN*, *CASQ2*, *RRAD, CAV2* and *S100A4,* metabolism-related *MFN2*, *SLC6A8,* and *GPX3* were consistently upregulated in HPLM/B27-treated cells compared to RPMI/B27_glcM (Figures 2B-C).

PCA of qPCR data resolved three distinct clusters with HPLM/B27-treated hPSC-CMs occupying a gene expression space distinct from RPMI/B27_glcM and Day 16 groups (Figure 2D). This separation was driven by genes critical to structural and functional maturation, including upregulated *MYL2*, *CSRP3*, *PLN*, and downregulated *MYH6* (Figure 2E). These results underscore the role of HPLM in mediating the expression of sarcomere and calcium handling genes related to hPSC-CM maturity (Figure 2E).

In summary, transcriptomic analyses revealed that HPLM/B27 enhanced hPSC-CM maturation beyond that achieved with reduced glucose RPMI formulations alone. This suggests that the differential availability of other defined components found in HPLM versus RPMI promotes a coordinated maturation program that spans structural, functional, and metabolic phenotypes.

### 3.2 HPLM promotes structural maturation of hPSC-CMs

To determine whether the transcriptomic changes observed in HPLM-cultured hPSC-CMs translate to enhanced structural maturation, we assessed sarcomere organization and structural features of Day 30 hPSC-CMs cultured in HPLM/B27 versus RPMI_glcM/B27 to focus on effects of HPLM beyond reduced glucose concentrations.

Building on transcriptomic findings that identified changes in sarcomere isoform expression in HPLM/B27, we compared Day 30 hPSC-CMs cultured in HPLM/B27 versus RPMI_glcM/B27 using immunostaining (Figure 3B-C; 3E-F). RNA-seq and qPCR analyses revealed a significant increase in the *MYH7*/*MYH6* and *MYL2*/*MYL7* isoform ratio (Figure 3A and 3D), indicative of a shift toward mature ventricular-like myosin expression. Consistent with this, immunostaining for β-MHC and α-MHC indicated a predominant expression of β-MHC. MLC2v and MLC2a immunostaining demonstrated a marked increase in the proportion of MLC2v^+^/MLC2a^-^ cells and a corresponding decrease in the MLC2v^-^/MLC2a^+^ population in HPLM/B27-treated cells (Figures 3E–F), confirming an accelerated myosin light chain isoform switch at the protein level. Although RNA-seq analysis identified a higher *TNNI3*/*TNNI1* ratio in one differentiation (Figure 3G), qPCR validation across independent differentiations failed to confirm this difference. This suggests that later-stage sarcomere maturation had not yet been achieved during the 2-week culture period in HPLM/B27.

**Figure 3.**
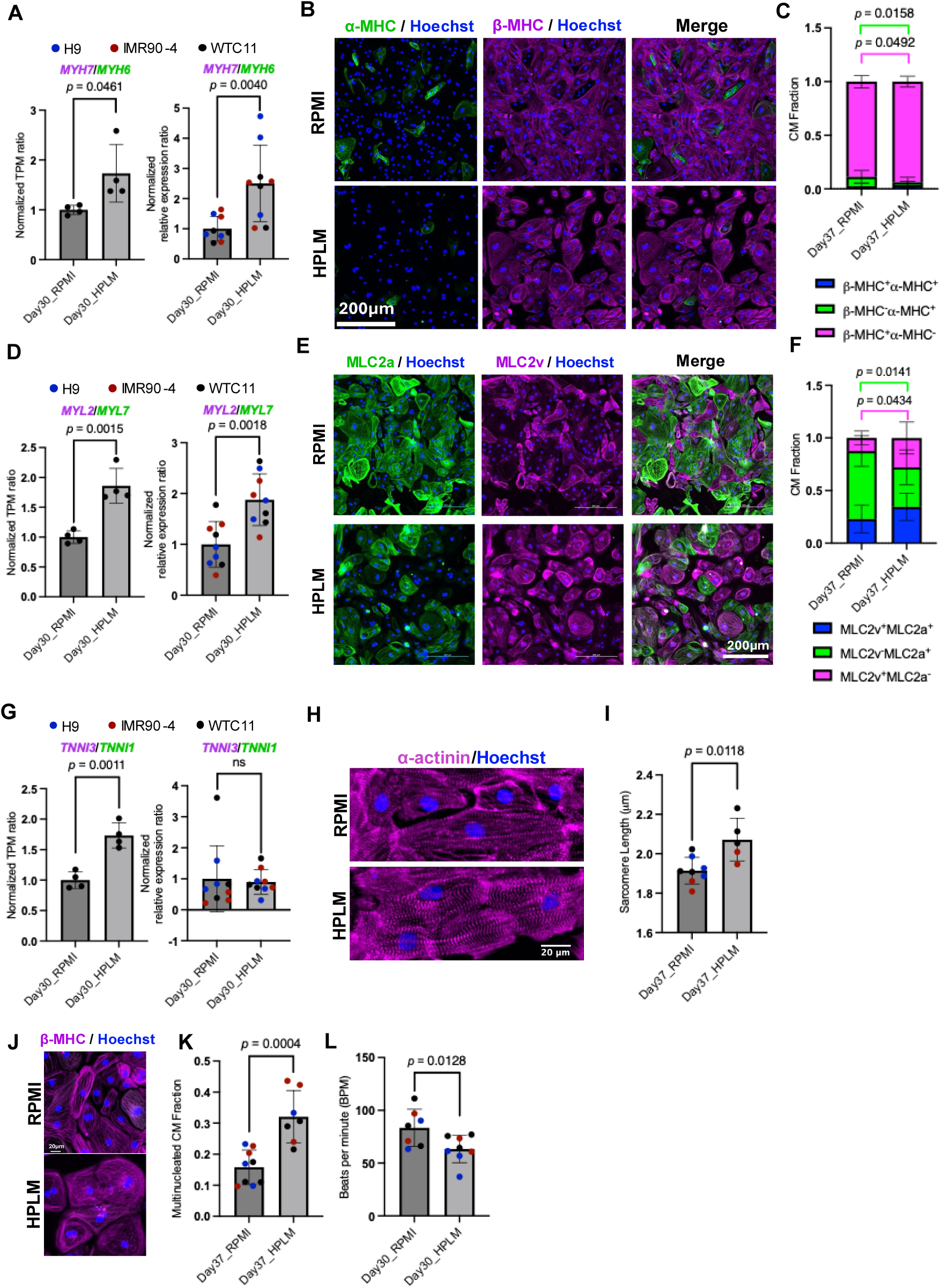
HPLM promotes structural maturation of hPSC-CMs. (A) Relative expression ratios of *MYH7* to *MYH6* by RNA-seq (left, n = 4) and qPCR (right, n = 9), comparing Day 30 hPSC-CMs cultured in RPMI_glcM/B27 (RPMI/B27) vs. HPLM/B27. Data points are color-coded by cell line (blue: H9, red: IMR90-4, black: WTC11). (B) Representative immunostaining images for β-MHC (magenta), α-MHC (green), and Hoechst nuclear (blue) stains in Day 37 hPSC-CMs cultured in (upper) RPMI_glcM/B27 and (lower) HPLM/B27. Day 30 cells cultured in the indicated medium were replated on Matrigel-coated ibidi 96-well plates and maintained in the same medium for 7 days before immunofluorescent staining. Scale bar = 200 μm. (C) Quantification of β-MHC and α-MHC expression in Day 37 hPSC-CMs cultured in HPLM/B27 or RPMI_glcM/B27. Blinded immunostaining images were visually analyzed to determine the fraction of cells expressing β-MHC only (single-positive), α-MHC only (single-positive), or both (double-positive). For each condition, two random fields per image were counted (≥20 cells per image), with each image derived from an independent differentiation. Data were pooled from three hPSC lines (H9, IMR90-4, WTC11) across independent experiments. Sample sizes: RPMI_glcM/B27 (n=9 images), HPLM/B27 (n=7 images). Each dot represents one image from one independent differentiation. (D) Relative expression ratios of *MYL2* to *MYL7* by RNA-seq (left, n = 4) and qPCR (right, n = 9), comparing Day 30 hPSC-CMs cultured in RPMI_glcM/B27 (RPMI/B27) vs. HPLM/B27. Data points are color-coded by cell line (blue: H9, red: IMR90-4, black: WTC11). (E) Representative immunostaining images for MLC2v (magenta), MLC2a (green), and Hoechst nuclear (blue) stains in Day 37 hPSC-CMs cultured in (upper) RPMI_glcM/B27 and (lower) HPLM/B27. Day 30 cells cultured in the indicated medium were replated on Matrigel-coated ibidi 96-well plates and maintained in the same medium for 7 days prior to immunofluorescent staining. Scale bar = 200 μm. (F) Quantification of MLC2v and MLC2a expression in Day 37 hPSC-CMs cultured in HPLM/B27 or RPMI_glcM/B27. Blinded immunostaining images were visually analyzed to determine the fraction of cells expressing MLC2v only (single-positive), MLC2a only (single-positive), or both (double-positive). For each condition, two random fields per image were counted (≥20 cells per image), with each image derived from an independent differentiation. Data were pooled from three hPSC lines (H9, IMR90-4, WTC11) across independent experiments. Sample sizes: RPMI_glcM/B27 (n=9 images), HPLM/B27 (n=7 images). Each dot represents one image from one independent differentiation. (G) Relative expression ratios of *TNNI3* to *TNNI1* by RNA-seq (left, n = 4) and qPCR (right, n = 9), comparing Day 30 hPSC-CMs cultured in RPMI_glcM/B27 (RPMI/B27) vs. HPLM/B27. Data points are color-coded by cell line (blue: H9, red: IMR90-4, black: WTC11). (H) Representative immunostaining images with sarcomeric alpha-actinin (magenta), and Hoechst nuclear (blue) stains in Day 37 hPSC-CMs cultured in (upper) RPMI_glcM/B27 and (lower) HPLM/B27. Day 30 cells cultured in the indicated medium were replated on Matrigel-coated ibidi 96-well plates and maintained in the same medium for 7 days prior to immunofluorescent staining. Scale bar = 20 μm. (I) Quantification of sarcomere length in Day 37 hPSC-CMs cultured in RPMI_glcM/B27 and HPLM/B27. Sarcomere length was quantified from sarcomeric α-actinin immunofluorescence images (40x magnification; 6.25 pixels/µm) using SotaTool, For each condition, one image per independent differentiation (spanning three hPSC lines: H9, IMR90-4, WTC11) was analyzed with the following parameters: minimum sarcomere length = 1.0 µm, maximum sarcomere length = 2.5 µm, 4×4 grid segmentation, offset = 4 µm, and default settings for remaining parameters. Sample sizes: RPMI_glcM/B27 (n=8 images), HPLM/B27 (n=5 images). Each dot represents one image from one independent differentiation. (J) Representative immunostaining images for β-MHC (magenta) and Hoechst nuclear (blue) stains in Day 37 hPSC-CMs cultured in (upper) RPMI_glcM/B27 and (lower) HPLM/B27. Scale bar = 20 μm. (K) Quantification of multinucleation in Day 37 hPSC-CMs cultured in RPMI_glcM/B27 and HPLM/B27 on images stained with β-MHC and Hoechst. Manual counts of mononuclear and multinuclear cells were performed on blinded images (two random fields per image; ≥40 cells per image), with each image derived from an independent differentiation. Data were pooled from three hPSC lines (Blue: H9, Red: IMR90-4, Black: WTC11) across independent experiments. Sample sizes: RPMI_glcM/B27 (n=9 images), HPLM/B27 (n=7 images). Each dot represents one image from one independent differentiation. (L) Quantification of spontaneous beating rate for Day 30 hPSC-CMs cultured in RPMI_glcM/B27 and HPLM/B27 by using MUSCLEMOTION. Each dot represents the beat rate from a single video recording (50 frames per second, 20 seconds, 20x magnification) of one well, quantified using MUSCLEMOTION software, with each video derived from an independent differentiation. Data are pooled from three hPSC lines (Blue: H9, Red: IMR90-4, Black: WTC11) across independent experiments. Sample sizes: RPMI_glcM/B27 (n=7 videos), HPLM/B27 (n=8 videos). Data are mean ± SD, with the p-values shown in each panel determined by Student’s t-test.

Upregulation of *CSRP3* (cardiac LIM protein/MLP) in HPLM/B27-treated hPSC-CMs (Figure 2E) is consistent with its role as a regulator of sarcomere organization^51–54^. In line with these findings, immunostaining analysis revealed elongated sarcomeres (1.91 ± 0.02 µm to 2.07 ± 0.04 µm; p < 0.01; Figures 3H–I) in HPLM/B27-treated cells, further reflecting sarcomere maturation. This structural remodeling was accompanied by a doubling in multinucleation frequency (15.8 ± 1.7% to 32.1 ± 2.9%; p < 0.001; Figures 3J–K) in HPLM/B27-treated cells, a feature associated with cardiomyocyte maturity. HPLM/B27-treated cells exhibited significantly reduced spontaneous contraction rates (from 83 ± 6 BPM to 63 ± 4 BPM; Figure 3L; Videos S1 and S2), aligning with prior observations that decreased automaticity correlates with structural and electrophysiological maturation *in vitro*^55^.

Collectively, these findings demonstrate that HPLM/B27 promoted structural maturation in hPSC-CMs, characterized by myosin isoform switching, elongated sarcomeres, increased multinucleation, and reduced spontaneous beating rates.

### 3.3 HPLM enhances the Ca^2+^ handling capacity of Day 30 hPSC-CMs

To investigate whether HPLM-induced transcriptomic changes in calcium-handling genes translate to functional improvements in calcium transient kinetics, we performed real-time intracellular calcium imaging in Day 30 hPSC-CMs cultured in HPLM/B27 versus RPMI_glcM/B27 to focus on effects of HPLM beyond reduced glucose concentrations.

During maturation, cardiomyocytes develop more efficient calcium cycling dynamics with larger calcium transient amplitudes, faster calcium release from the SR, and accelerated calcium reuptake. RNA-seq and qPCR data revealed that HPLM/B27 significantly upregulated key calcium-handling genes compared to RPMI_glcM/B27, including *PLN*, *CASQ2*, *S100A4*, *ATP2A2*, and *ATP2A3* (Figure 1E and Figure 2B). Analysis of the calcium transient traces of Day 30 hPSC-CMs after a two-week culture in RPMI_glcM/B27 or HPLM/B27 confirmed that HPLM/B27 treatment led to approximately a 50% increase in peak amplitude compared to RPMI_glcM/B27 (from 0.756 ± 0.36 to 1.15 ± 0.13 ΔF/F_0_) (Figures 4A and 4B, Videos S3 and S4). Furthermore, HPLM/B27 significantly increased both the upstroke velocity (from 2.23 ± 0.26 to 4.17 ± 0.72 ΔF/F_0_/s) (Figure 4C) and downstroke velocity (from 1.23 ± 0.18 to 1.94 ± 0.22 ΔF/F_0_/s) (Figure 4D), consistent with more rapid calcium release and reuptake.

**Figure 4.**
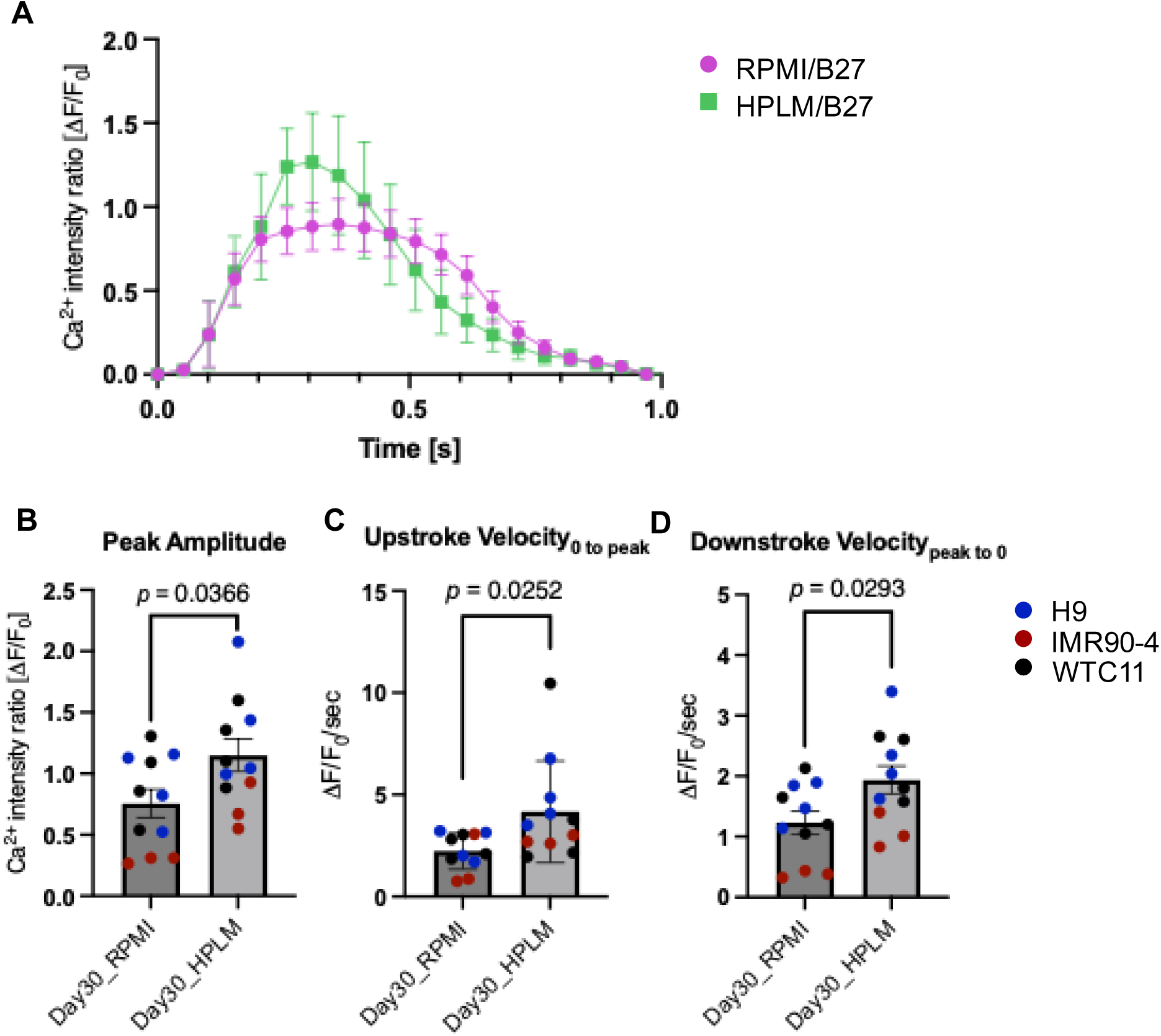
HPLM enhances Ca^2+^ cycling properties of hPSC-CMs. (A) Representative Ca^2+^ transients of hPSC-CMs cultured in HPLM/B27 and RPMI_glcM/B27. Cells were loaded with FLIPR Calcium 6 dye and fluorescence was recorded in real time at 20x magnification. Each trace shows a normalized fluorescence (ΔF/F₀) peak averaged from 20 cells per well, with error bars representing the mean ± SD across 3 wells per condition. (B-D) Quantification of Ca^2+^ handling (B) peak amplitude (ΔF/F₀), (C) upstroke velocity (ΔF/F/sec, baseline to peak), and (D) downstroke velocity (ΔF/F/sec, peak to baseline) from recorded Ca^2+^ transients in Day 30 hPSC-CMs cultured in HPLM/B27 and RPMI_glcM/B27. Each symbol represents the mean of 20 cells in one well, in H9 (blue, n=4 wells), IMR90-4 (red, n=3 wells), and WTC11 (black, n=4 wells) hPSCs. Data are presented as mean ± SD; p-values were calculated by Student’s t-test.

These results demonstrate that HPLM/B27 improved calcium handling in hPSC-CMs compared to RPMI_glcM/B27, as evidenced by increased calcium transient amplitudes and accelerated calcium release and reuptake kinetics, which more closely resemble mature cardiomyocyte function.

### 3.4 HPLM promotes oxidative and glycolytic metabolism of Day 30 hPSC-CMs

To determine whether HPLM-induced transcriptomic changes in metabolic genes translate to functional metabolic maturation, we performed bioenergetic profiling of Day 30 hPSC-CMs cultured in HPLM/B27 versus RPMI_glcM/B27 using Seahorse metabolic flux analysis, mitochondrial DNA copy number quantification, and fluorescence lifetime imaging of NAD(P)H.

Metabolic maturation is a hallmark of cardiomyocyte development, marked by a shift from glycolytic dependence to oxidative phosphorylation (OXPHOS) and fatty acid oxidation (FAO) to meet the energetic demands of continuous contraction^58,59^. RNA-seq analysis revealed that HPLM/B27-treated hPSC-CMs demonstrated upregulated biological processes associated with mitochondrial metabolism and glycolysis, alongside increased expression of metabolic genes critical for OXPHOS (*COX7B*, *SDHB*, *SDHD*), creatine metabolism (*SLC6A8*, *CKM*), and lipid metabolism (*LPL*, *PPARA*, *ACADVL*, *PPARGC1A*), consistent with more mature metabolism compared to hPSC-CMs cultured in RPMI_glcM/B27 (Figure 1D–E; Figure S2B).

To determine whether these changes in gene expression led to altered metabolic pathway utilization, we first performed the Seahorse Cell Mito Stress Test, which uses mitochondrial pathway inhibitors to measure oxygen consumption rate (OCR) as the primary output for oxidative metabolism, and extracellular acidification rate (ECAR) was concurrently measured as a secondary readout of glycolytic activity. To isolate glycolysis-specific effects, we focused on basal ECAR (Figure 5A–F). Consistent with the transcriptomic data, hPSC-CMs treated with HPLM/B27 exhibited 52% higher basal (from 45 ± 16 to 69 ± 58 pmol/min/10k cells) and 36% higher maximal (from 133 ± 49 to 180 ± 115 pmol/min/10k cells) OCR compared to RPMI_glcM/B27-treated cells (Figure 5C-D), indicating enhanced oxidative activity. Basal ECAR was 38% increased (from 37 ± 13 to 51 ± 31 mpH/min/10k cells) in HPLM/B27-treated cells (Figure 5E), suggesting transient co-activation of glycolysis in these cells. Notably, the OCR/ECAR ratio, a marker of metabolic preference^60^, remained unchanged between conditions (Figure 5F), demonstrating that both oxidative and glycolytic metabolism increased proportionally and suggesting a shift from glycolysis to oxidative phosphorylation is ongoing and not yet fully established at this timepoint.

**Figure 5.**
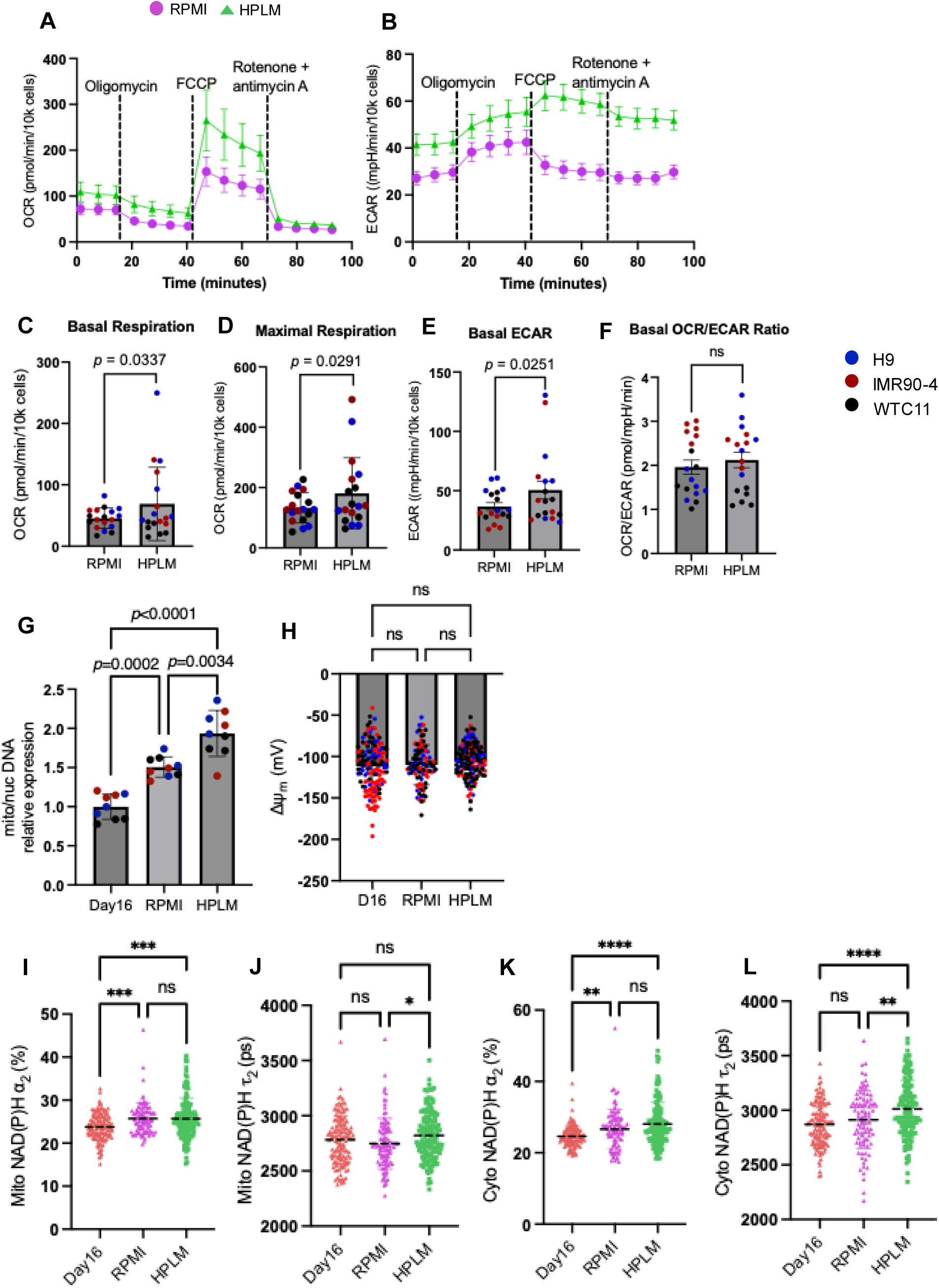
HPLM enhances oxidative and glycolytic metabolism of hPSC-CMs. (A-F) Mito stress test on Day 30 hPSC-CMs cultured in HPLM/B27 and RPMI_glcM/B27. Representative traces of real-time (A) oxygen consumption rate (OCR) and (B) extracellular acidification rate (ECAR) in hPSC-CMs cultured in HPLM/B27 (green) or RPMI_glcM/B27 (purple) measured by Seahorse extracellular flux analyzer. Cells were treated with Oligomycin (ATP synthase inhibitor), FCCP (respiratory uncoupler), and Rotenone and antimycin A (respiratory chain blockers). C-F. Statistical analysis of (C) basal respiration (D) maximal respiration (E) basal ECAR before adding oligomycin and (F) ratio of basal OCR and ECAR comparing Day 30 hPSC-CMs cultured in HPLM/B27 and RPMI_glcM/B27. Data include hPSC-CMs derived from H9 (blue), IMR90-4 (red), and WTC11 (black) hPSCs, with n=3 differentiation batches per cell line. Data are presented as mean ± SD; p-values are calculated by Student’s t-test. (G) Ratio of mitochondrial to nuclear DNA content in Day 16 hPSC-CMs cultured in RPMI/B27, and Day 30 hPSC-CMs cultured in RPMI_glcM/B27 or HPLM/B27. Relative expression levels were normalized to the Day 16 condition for both HPLM and RPMI conditions. Data include hPSC-CMs derived from H9 (blue), IMR90-4 (red), and WTC11 (black) hPSCs, with n = 3 independent differentiation batches per cell line. Data are presented as mean ± SD, and statistical significance is determined by one-way ANOVA with Tukey’s post hoc test. (H) Single-cell mitochondrial membrane potential (MMP) measured by TMRE in Day 16 hPSC-CMs cultured in RPMI/B27, and Day 30 hPSC-CMs cultured in RPMI_glcM/B27 or HPLM/B27. MMP (mV) was quantified by fluorescence intensity of TMRE (FI_m_) and nuclear background fluorescence intensity (FI_n_) based on Nernst Equation^56,57^. Each dot represents a single cell, and data include three independent differentiation batches from H9 (blue), IMR90-4 (red), and WTC11 (black) hPSCs. Data are presented as mean ± SD, and statistical significance is determined by one-way ANOVA with Tukey’s post hoc test. (I-L) Single-cell (I-J) mitochondrial and (K-L) cytoplasmic NAD(P)H autofluorescence lifetime parameters (I) Mitochondrial NAD(P)H protein-bound fraction (J) Mitochondrial fluorescence lifetime of the bound form of NAD(P)H. (K) Cytoplasmic NAD(P)H protein-bound fraction (L) Cytoplasmic fluorescence lifetime of the bound form of NAD(P)H of hPSC-CMs in Day 16 hPSC-CMs cultured in RPMI/B27, and Day 30 hPSC-CMs cultured in RPMI_glcM/B27 or HPLM/B27. Data include 90 to 130 single cells for three independent differentiation batches of H9, IMR90-4, and WTC11 hPSCs. Data are presented as mean ± SD and statistical significance is determined by one-way ANOVA with Tukey’s post hoc test. *p<0.05, **p<0.01, ***p<0.001, ****p<0.0001

We next investigated mitochondrial biogenesis and function, as HPLM/B27 also upregulated mitochondrial-related GO terms (Figure 1D) and increased expression of *MFN2* (mitofusin 2), a key regulator of mitochondrial fusion (Figures 2C and 2E). qPCR analysis revealed a 0.28-fold increased mitochondrial-to-nuclear DNA ratio in HPLM/B27-treated cells compared to RPMI_glcM/B27 (Figure 5G), indicating an increased total mitochondrial content. However, mitochondrial membrane potential, measured by TMRE intensity, remained unchanged across different conditions (Figure 5H), suggesting that HPLM/B27 enhanced metabolic capacity at this time point primarily through mitochondrial biogenesis rather than hyperpolarizing individual mitochondria.

To more deeply probe the enhancement of oxidative and glycolytic metabolism observed in HPLM/B27-treated hPSC-CMs, we performed fluorescence lifetime imaging (FLIM) of NAD(P)H autofluorescence (Figure S3) on Day 30 hPSC-CMs treated by RPMI_glcM/B27 and HPLM/B27. NAD(P)H serves as an endogenous fluorescent metabolic coenzyme in both glycolytic and oxidative pathways, with its protein-binding characteristics, particularly binding portion (α2) and duration (t2), providing insights into metabolic enzyme engagement and pathway efficiency. We spatially resolved NAD(P)H dynamics in mitochondrial and cytoplasmic compartments via two-photon microscopy by labeling mitochondria with TMRE (Figure 5I–L; Figure S3). Mitochondrial analysis revealed no significant difference in the protein-bound NAD(P)H fraction (α2) between HPLM/B27 and RPMI_glcM/B27 (Figure 5I); however, both Day 32 groups exhibited elevated α2 compared to Day 16 baseline differentiation, indicating metabolic maturation-associated increases in mitochondrial NAD(P)H engagement with oxidative enzymes. Interestingly, HPLM/B27-treated cells showed prolonged NAD(P)H-protein binding times (t2) in mitochondria compared to RPMI_glcM/B27 (Figure 5J), indicating changes in preferred NAD(P)H binding partners within the mitochondria that could result in the observed elevated basal respiration (Figures 5A and 5C). Similarly, cytoplasmic NAD(P)H analysis mirrored mitochondrial trends, with α2 significantly increased in Day 32 groups versus Day 16 (Figure 5K) and marginally prolonged t2 (Figure 5L) in HPLM/B27-treated cells compared to RPMI_glc/B27, consistent with transcriptomic upregulation of glycolytic pathways (Figure S2B) and elevated basal ECAR (Figure 5E). These findings demonstrate that HPLM/B27 drove coordinated metabolic remodeling with prolonged NAD(P)H-enzyme interactions in mitochondria that enhance oxidative metabolism, with glycolytic activation occurring predominantly in the cytoplasm.

Collectively, these findings demonstrate that HPLM/B27 promotes metabolic maturation in hPSC-CMs through coordinated bioenergetic remodeling, characterized by increased mitochondrial biogenesis, enhanced OXPHOS efficiency via prolonged NAD(P)H-enzyme interactions, and transient glycolytic co-activation. While the metabolic shift toward oxidative metabolism appears to be in progress rather than complete after two weeks of culture, HPLM/B27 accelerates this transition compared to RPMI_glcM/B27.

## 4. Discussion

Structural maturation of cardiomyocytes during human cardiac development includes sarcomere protein isoform transitions. As the GiWi protocol mainly produces ventricular-CMs^34,35,62^, we evaluated the expression of ventricular sarcomere maturation markers (*MYH7* and *MYL2*). Our immunostaining results confirmed the predominant expression of the matured myosin heavy chain isoform (β-MHC) in HPLM/B27 conditions (Figure 3B-C), highlighting that this early maturation baseline was achieved. Next, quantitative gene expression analysis demonstrated a progressive increase in the *MYL2*/*MYL7* ratio (Figure 3A), with HPLM/B27 cultures showing a significantly higher MLC2v+ population by immunostaining (Figure 3B-C). This aligns with prior work reporting comparable MLC2v+ populations in Day 27 hPSC-CMs following one week of 3D engineered cardiac patch maturation ^63^. Together, these findings suggest that while physiologic glucose suffices for initiating the myosin heavy chain isoform switch (α-MHC to β-MHC), HPLM/B27 accelerates the later-stage myosin light chain transition (MLC2a to MLC2v) by Day 30. Additionally, *CSRP3*, which encodes Muscle LIM Protein (MLP), a scaffold for sarcomere assembly, was markedly upregulated in HPLM/B27-treated cells compared to the physiologic glucose RPMI/B27 group (Figure 2E). Although *CSRP3*’s role in hPSC-CM maturation remains understudied, *CSRP3* deficiency disrupts Z-disc integrity, underscoring its importance for sarcomere organization ^64,65^. Supporting this, we observed elongated sarcomere length (∼2.07 µm) after HPLM/B27 treatment, approaching the adult human lengths achieved by other maturation protocols *in vitro* ^7,20,66^.

Given the direct correlation between structural maturity and calcium-handling capacity in cardiomyocytes^7^, we evaluated functional maturity in Day 30 hPSC-CMs by measuring intracellular calcium transients in HPLM/B27 versus physiologic glucose RPMI/B27 cultures. Transcriptomic analysis revealed significant upregulation of calcium-handling machinery genes including *SLC8A1*, *KCNJ2*, *ATP2A2*, *CASQ2*, *PLN*, and *S100A4* in HPLM/B27-treated cells. Consistent with these changes in gene expression, HPLM/B27-cultured hPSC-CMs exhibited increased calcium transient peak amplitudes, along with faster upstroke and downstroke velocities, indicative of enhanced cytosolic calcium dynamics and improved sarcoplasmic reticulum calcium reuptake/release efficiency. These findings align with prior studies demonstrating that optimized metabolic maturation media enhance calcium-handling properties in hPSC-CMs^24,25,67–69^, suggesting a potential metabolism-dependent mechanism underlying this functional improvement. Notably, while HPLM provides roughly six-fold more calcium (2.4 mM) than RPMI (0.42 mM), this difference alone cannot likely explain the observed enhancement of calcium-handling capacity. A previous study reported reduced calcium transient amplitudes under electrical pacing for hPSC-CMs cultured in conditions with 1.8 mM versus 0.42 mM calcium ^70^, highlighting the nuanced role of calcium concentration. This implies that HPLM’s benefits likely arise from a synergistic interplay of physiological ions and polar metabolites rather than elevated calcium alone.

For metabolic maturation, HPLM/B27-treated hPSC-CMs exhibited elevated mitochondrial content alongside higher basal and maximal oxidative respiration, aligning with earlier reports ^23,24,67,68,71^. Intriguingly, despite adult cardiomyocytes relying predominantly on oxidative metabolism with reduced glycolysis ^7,18,72^, increased basal ECAR was also observed in HPLM/B27 cultures. Consistent with our finding, other studies also observed increased OCR and ECAR in matured hPSC-CMs when supplementing oxidative substrates ^24,68^ which suggests that hPSC-CMs might demonstrate metabolic flexibility and can switch to glucose utilization during the turnover of metabolic maturation while fatty acid oxidation related machinery is not fully established^68,73^. To further validate the observed increases in oxidative and glycolytic activity, we applied fluorescence lifetime imaging microscopy (FLIM) to measure NAD(P)H lifetime at single-cell resolution, with specific attention to mitochondrial and cytoplasmic compartments and found consistently prolonged NAD(P)H lifetimes (t2) in both mitochondrial and cytoplasmic compartments, while the bound fraction (a2) remained unchanged compared to physiologic glucose RPMI/B27. NAD(P)H is a central cofactor in mitochondrial function and energy metabolism, and its lifetime dynamics reflect metabolic activity ^74,75^. Given the spatial compartmentalization of metabolic pathways where glycolysis predominantly occurs in the cytoplasm, while oxidative metabolism is localized to mitochondria ^76^, FLIM provides distinct spatial resolution at the single-cell level, overcoming limitations of invasive bulk measurements that average signals across heterogeneous cell population ^77,78^.

While high glucose levels inhibit *in vitro* hPSC-CM maturation^79^, achieving an optimal balance between glucose concentration and other carbon sources appears critical for advancing maturation ^24,79,80^. Notably, our transcriptomic analyses revealed that *MYL2* (Figure 1E) was expressed at lower levels in physiologic glucose RPMI/B27 cultures compared to RPMI/B27. This observation suggests that while physiological glucose levels in HPLM may initiate metabolic maturation, additional carbon sources in HPLM (e.g., lactate and galactose) likely synergize to meet the energetic demands of enhanced maturation. This hypothesis aligns with prior studies in which cardiac maturation media, modified from RPMI/DMEM through glucose reduction and supplementation with lactate, galactose, or fatty acids, improved hPSC-CM structural, functional, and metabolic maturity ^23,24^. Similarly, HPLM provides several additional metabolites, which may collectively explain its enhanced maturation outcomes compared to physiologic glucose RPMI/B27.

It is notable that HPLM contains metabolites previously associated with hPSC-CM maturation. For example, HPLM includes both glutathione, a key mediator of redox homeostasis, and taurine, a metabolite that prior studies have supplemented to improve hPSC-CM maturation^24,81^. This combination suggests potential synergy in antioxidant pathways, as glutathione and taurine are both critical for redox balance. Consistent with this, our RNA-seq and qPCR analyses revealed elevated *GPX3* expression in HPLM/B27-treated hPSC-CMs, indicative of enhanced antioxidant activity. While fatty acids were intentionally excluded to evaluate HPLM as a standalone basal medium, HPLM contains acetylcarnitine and carnitine, metabolites critical for fatty acid transport and oxidation. Carnitine has been proposed to enhance fatty acid metabolism during hPSC-CM maturation ^24,67^. Supporting this, we observed upregulated expression of genes linked to fatty acid β-oxidation and mitochondrial regulation (*ACADVL* and *PPARGC1A*) in HPLM/B27 cultures. Furthermore, HPLM incorporates ketone bodies such as 3-hydroxybutyrate, metabolites increasingly recognized as important drivers of cardiomyocyte maturation ^82^. By recapitulating the ionic and polar metabolite composition of adult human plasma, HPLM establishes a physiologically balanced basal environment that may intrinsically support critical maturation pathways. This positions HPLM as a robust platform for future studies aiming to optimize hPSC-CM maturation through targeted supplementation or combinatorial approaches.

Despite its promise, our study has limitations that warrant further investigation. While comparing HPLM/B27 to RPMI/B27 allowed us to identify benefits of a physiologic medium designed based on adult plasm, it precluded the inclusion of supplementary factors previously implicated in hPSC-CM maturation, such as fatty acids, glucocorticoids, triiodothyronine (T3), insulin-like growth factors (IGFs), and peroxisome proliferator-activated receptor (PPAR) agonists^1,25–27,69,83^. Beyond biochemical supplementation, integrating biophysical cues reflective of the adult cardiac microenvironment, such as mechanical loading ^21^, electrical stimulation ^19^, or co-culture with non-myocytes ^22^, may further advance functional maturity. Also, extending HPLM to disease-specific or patient-derived hPSC-CM models would clarify its utility in modeling cardiac pathologies or examining cardiotoxicity. Collectively, these efforts including refining media composition through additive strategies and diversifying experimental models could position HPLM as a versatile platform for advancing hPSC-CM maturation and translational applications.

## 5. Conclusion

hPSC-CMs cultured for two weeks in HPLM/B27 exhibited increased structural, functional, and metabolic maturation compared to cells cultured in RPMI/B27. Transcriptomic analysis indicated that the maturation benefit of HPLM surpassed that of physiologic glucose treatment in RPMI 1640 basal medium. Structurally, HPLM increased MLC2v^+^ population, elongated sarcomere length, and increased multinucleation percentage, accompanied by slower spontaneous beating rates. Calcium handling properties were markedly enhanced in HPLM cultures, as evidenced by larger calcium transient amplitudes, faster kinetics, and upregulated expression of calcium regulatory genes. Metabolic profiling demonstrated improvements in basal oxidative capacity and mitochondrial content, alongside elevated glycolytic activity. Collectively, these findings highlight HPLM/B27 as a versatile culture tool that advances hPSC-CM maturation, enabling more robust in vitro modeling of human cardiac physiology.

## Data availability

RNAseq: GSE294004. Key resources and HPLM medium formulation are detailed in Table S1 and S3 respectively.

## Acknowledgements

This work was supported by the National Science Foundation Center for Cell Manufacturing Technologies (CMaT, grant EEC-1648035) and the National Institutes of Health (grants R01HL165726, R01HL148059, R01HL178095, K22CA225864, R35GM156513, T32GM008349; T32GM140935, T32GM135066, T32HG002760, and F30HL173988). The authors thank Kayvan Samimi, Wenxuan Zhao, and Ty Weaver for their assistance and input on this project.

