## Supporting Information for "Human Plasma-Like Medium Promotes Maturation of Human Pluripotent Stem Cell-Derived Cardiomyocytes"

**Table S1. Reagents and resources used in this study**

| REAGENT or RESOURCE | SOURCE | IDENTIFIER |
|---|---|---|
| **Antibodies** | | |
| rb-IgG-anti-MYH7 | R&D Systems | Cat#MAB90961100 |
| ms-IgG1-anti-MYH6 | R&D Systems | Cat#MAB8979 |
| ms-IgG1 anti-cTnT | ThermoFisher | Ca#MA5–12960 |
| Alexa Fluor 488-conjugated anti-mouse IgG1 | ThermoFisher | Cat# A-21121 |
| mouse IgG2b anti-MLC2A | Synaptic Systems | Cat #311011 |
| rabbit IgG anti-MLC2V | Proteintech | Cat#10906-1-AP |
| Alexa Fluor 488 anti-mouse IgG2b | ThermoFisher | Cat# A11008 |
| Alexa Fluor 647 anti-rabbit IgG | ThermoFisher | Cat# A21240 |
| mouse IgG1 anti-α-actinin | Sigma | Cat# A7811 |
| Alexa Fluor 488 anti-mouse IgG2b | ThermoFisher | Cat#A21141 |
| Alexa Fluor 647 anti-mouse IgG1 | ThermoFisher | Cat#A21240 |
| **Chemicals, peptides, and recombinant proteins** | | |
| Growth Factor Reduced Matrigel | Corning | Cat#354263 |
| Versene | Life Technologies | Cat#15040066 |
| Accutase | Innovative Cell Technology | Cat#AT104 |
| Y-27632 | Tocris | Cat#1254 |
| CHIR99021 | Selleckchem | Cat#S1263 |
| IWP2 | Tocris | Cat#3533 |
| Fetal bovine serum | R&D Systems | Cat#S12450 |
| DMSO | Sigma-Aldrich | Cat#D2650 |
| Paraformaldehyde | Electron Microscopy Sciences | Cat#15710-S |
| Bovine Serum Albumin | ThermoFisher | Cat#BP1600 |
| Triton X-100 | ThermoFisher | Cat#BP151 |
| Hoechst 33342 | Invitrogen | Cat#H3570 |

| | | |
|---|---|---|
| Trizol Reagent | ThermoFisher | Cat#15596018 |
| RNaseOUT Recombinant Ribonuclease Inhibitor | Life Technologies | Cat#10777-019 |
| Oligo dT(20) primers | Life Technologies | Cat#18418020 |
| PowerUp SYBR Green Master Mix for qPCR | ThermoFisher | Cat#25780 |
| **Experimental models: Cell lines** | | |
| Human: IMR90-4 hiPSC line | N/A | N/A |
| Human: WTC11 hiPSC line | N/A | N/A |
| Human: H9 hESC line | N/A | N/A |
| **Software and algorithms** | | |
| MUSCLEMOTION (version 1.1) | N/A | https://github.com/l-sala/MUSCLEMOTION |
| Sotatool | N/A | https://github.com/steinjm/SotaTool |
| ImageJ | N/A | https://imagej.nih.gov/ij/ |
| GraphPad Prism (version 9.4.1) | N/A | https://graphpad.com |
| BioRender | N/A | https://biorender.com |
| Nikon Instruments Software Elements (version 5.30.06) | N/A | https://www.microscope.healthcare.nikon.com/products/software/nis-elements |
| MetaboAnalyst (version 6.0) | N/A | https://www.metaboanalyst.ca/MetaboAnalyst/ModuleView.xhtml |
| **Critical Commercial Assays** | | |
| Zymo RNA Clean & Concentrator-25 | Zymo Research | Cat#R1018 |
| Qiagen Omniscript RT Kit | Qiagen | Cat#205113 |
| Seahorse XF Cell Mito Stress Kit | Agilent Technologies | N/A |

| | | |
|---|---|---|
| FLIPR® Calcium 6 Assay Kits | Molecular Devices | Ca#RB190 |
| Sequencing libraries prep | Takara | Ca#634411 |
| RNA sequencing | Illumina NovaSeq6000 | N/A |
| **Other** | | |
| mTeSR1 medium | STEMCELL Technologies | Cat#85850 |
| DMEM/F12 medium | ThermoFisher | Cat#11330032 |
| Standard RPMI1640 | Life Technologies | Cat#11875119 |
| Glucose free RPMI 1640 | Life Technologies | Cat#11879020 |
| HPLM | Rossiter et al. 2021[1] | See supplemental table 3 |
| B27 minus insulin supplement | Life Technologies | Cat#0050129SA |
| B27 plus insulin supplement | Life Technologies | Cat#17504044 |
| Mr. Frosty Freezing Container | ThermoFisher | Cat#51000001 |
| DPBS | ThermoFisher | Cat#14190144 |

### Table S2. Primer Sequences for qPCR

| Target | FW Primer (5'-3') | RV Primer (5'-3') |
|---|---|---|
| *ZNF384* | AATCTGCAGTCCCACAGACG | ACTGTGTGCGTAGACAGGTG |
| *EDF1* | CCAAGCAGGCTATCTTAGCGG | GACCTTGCTGGATCACCTTG |
| *DDB1* | TCAACGGCATGATAGGGCTG | CGCTCGGTGTGAAAGGATCT |
| *TNNT2* | TTCACCAAAGATCTGCTCCTCGCT | TTATTACTGGTGTGGAGTGGGTGTGG |
| *MYH6* | AGCTCACCTACCAGACAGAGG | TTGCTTGGCACCAATGTCAC |
| *MYH7* | GAGGAGCAAGCCAACACCAA | CTCATTCAAGCCCTTCGTGC |
| *MYL7* | GGTGGTGAACAAGGATGAGTT | GTGACTTGTAGTCGATGTTCCC |
| *MYL2* | GCTGAAGGCTGATTACGTTCG | TCCAAGTTGCCAGTCACGTC |
| *TNNI1* | GCAAACTCTTGCTGAAGAGCC | GTGTTGTGGAGGCATTTGGC |
| *TNNI3* | TGCAGATGCCATGATGCAGG | CCCGGTTTTCCTTCTCGGTG |
| *SYNPO* | GGGATCGAGGCTCAGGACC | GGCTCACCCAGCCGTCTA |
| *RRAD* | CGACTCAGACGAGAGCGTTT | GATCATAGGTGTGCCCTGCT |
| *GPX3* | ATGGGCAATCCCCAGATGGAC | GACCGAATGGTGCAAGCTCT |
| *PCDH20* | TGGGAAGCCACCCAGAGAAT | TTTCGTCCAGCGTCAGCATC |
| *CSRP3* | CCACACAGGCAGACTTGACC | CTGTCGTGCTGTCAAGAGCC |
| *CAV2* | GCTGTCTGCACATCTGGATTTTA | AATCCTGGCTCAGTTGCAGG |
| *MFN2* | GCTCGCTGGTGACGTAGTGA | TGTCTCAGGTTGAGGTTGGC |
| *PLN* | ATCACAGCTGCCAAGGCTAC | TGACGTGCTTGTTGAGGCAT |
| *GJA5* | AGCACATGGCTAAGTGCCAG | TCGTACTTGCTCGGTGACCA |
| *CASQ2* | GACGACTTTCCTCTGCTCGTT | CAGCTCCTCAGCAGTTGGAA |
| *SLC6A8* | CTGGGAGGTGACCCTTTGTCT | TAAATGATGCCATCCAGGGCG |
| *ITGA5* | AGACTTTCTTGCAGCGGGAG | ATCCACAGTGGGACGCCATA |
| *TNC* | GGACTCCTGTACCCCTTCCC | TGCGTCTCAGGAACACAATCC |
| *HIF1A* | GCAGAATGCTCAGAGAAAGCGA | GCTGCATGATCGTCTGGCTG |
| *MEF2C* | TGACTGTGAGATTGCGCTGA | CTTCTTTCTCAACGTCTCCACG |
| *MRPS6* | CACAACAGAGGCGGGTATTTCT | CGAGTGGGACTGGGACAATC |
| *S100A4* | CTGACTGCTGTCATGGCGTG | CAGCTTCATCTGTCCTTTTCCCC |
| *OPTN* | GAAAGGCCCGGAGACTGTTG | TCCTTTCAAGGGCCTGACAC |
| *ACADVL* | TTGTCCACCCGGAGTTGAGT | TGCAGCAGAAACTGTTCATTGAC |
| *DECR1* | TGGAAGCCATGAGCAAGTCTC | GGCGACCACAGGGAATTCTG |
| *PPARGC1A* | TGAACTGAGGGACAGTGATTTC | CCCAAGGGTAGCTCAGTTTATC |
| *nucDNA* | CAACTTCATCCACGTTCACC | GAAGAGCCAAGGACAGGTAC |
| *mitoDNA* | CGAAAGGACAAGAGAAATAAGG | CTGTAAAGTTTTAAGTTTTATGCG |

### Table S3. Basal HPLM formulation

| | | | | |
|---|---|---|---|---|
| **Concentrated stocks of components were prepared and pooled as described below** | | | | |
| **All working concentrations in uM** | | | | |
| Components added relative to initial formulation described before[2] | | | | |
| | Concentration | Vendor | Product # | Stock solution pool (Storage) |
| ***Salts*** | | | | |
| CaCl2 | 2350 | Sigma-Aldrich | C5670 | 10x (-20°C) |
| KCl | 4100 | Sigma-Aldrich | P5405 | 10x (-20°C) |
| MgCl2 | 480 | Sigma-Aldrich | M8266 | 10x (-20°C) |
| MgSO4 | 350 | Sigma-Aldrich | M2643 | 10x (-20°C) |
| NaCl | 105000 | Sigma-Aldrich | S7653 | 10x (-20°C) |
| NaHCO3 | 24000 | Sigma-Aldrich | S5761 | 10x (-20°C) |
| Na2HPO4 | 870 | Sigma-Aldrich | S9390 | 10x (-20°C) |
| Ca(NO3)2 | 40 | Sigma-Aldrich | C1396 | 100x (-20°C) |
| NH4Cl | 40 | Sigma-Aldrich | A9434 | 100x (-20°C) |
| ***Metabolites*** | | | | |
| Histidine | 110 | Sigma-Aldrich | H5659 | 500x (-20°C) prepared in 0.1M HCl |
| Isoleucine | 70 | Sigma-Aldrich | I2752 | 500x (-20°C) prepared in 0.1M HCl |
| Leucine | 160 | Sigma-Aldrich | L8000 | 500x (-20°C) prepared in 0.1M HCl |
| Lysine | 200 | Sigma-Aldrich | L5626 | 500x (-20°C) prepared in 0.1M HCl |
| Methionine | 30 | Sigma-Aldrich | M9625 | 500x (-20°C) prepared in 0.1M HCl |
| Phenylalanine | 80 | Sigma-Aldrich | P2126 | 500x (-20°C) prepared in 0.1M HCl |
| Threonine | 140 | Sigma-Aldrich | T8625 | 500x (-20°C) prepared in 0.1M HCl |
| Tryptophan | 60 | Sigma-Aldrich | T0254 | 500x (-20°C) prepared in 0.1M HCl |
| Valine | 220 | Sigma-Aldrich | V0500 | 500x (-20°C) prepared in 0.1M HCl |
| Alanine | 430 | Sigma-Aldrich | A7627 | 500x (-20°C) |
| Arginine | 110 | Sigma-Aldrich | A5131 | 500x (-20°C) |
| Asparagine | 50 | Sigma-Aldrich | A0884 | 500x (-20°C) |

| | | | | |
|---|---|---|---|---|
| Cysteine | 40 | Sigma-Aldrich | C1276 | 500x (-20°C) |
| Glycine | 300 | Sigma-Aldrich | G7126 | 500x (-20°C) |
| Proline | 200 | Sigma-Aldrich | P0380 | 500x (-20°C) |
| Serine | 150 | Sigma-Aldrich | S4500 | 500x (-20°C) |
| Aspartate | 20 | Sigma-Aldrich | A9256 | 500x (-20°C) prepared in 1M HCl |
| Cystine | 100 | Sigma-Aldrich | C8755 | 500x (-20°C) prepared in 1M HCl |
| Glutamate | 80 | Sigma-Aldrich | G1251 | 500x (-20°C) prepared in 1M HCl |
| Tyrosine | 80 | Sigma-Aldrich | T3754 | 500x (-20°C) prepared in 1M HCl |
| Glutamine | 550 | Sigma-Aldrich | G3126 | 250x (-20°C) |
| 4-hydroxyproline | 20 | Sigma-Aldrich | H5534 | 500x (-20°C) |
| Acetylcarnitine | 5 | Sigma-Aldrich | A6706 | 500x (-20°C) |
| Acetylglycine | 90 | Sigma-Aldrich | A16300 | 500x (-20°C) |
| alpha-Aminobutyrate | 20 | Sigma-Aldrich | A2536 | 500x (-20°C) |
| Betaine | 70 | Sigma-Aldrich | 61962 | 500x (-20°C) |
| Carnitine | 40 | Sigma-Aldrich | C0283 | 500x (-20°C) |
| Citrulline | 40 | Sigma-Aldrich | C7629 | 500x (-20°C) |
| Ornithine | 70 | Sigma-Aldrich | O2375 | 500x (-20°C) |
| Taurine | 90 | Sigma-Aldrich | T0625 | 500x (-20°C) |
| 2-hydroxybutyrate | 50 | Sigma-Aldrich | 220116 | 250x (-20°C) |
| 3-hydroxybutyrate | 50 | Sigma-Aldrich | 298360 | 250x (-20°C) |
| Acetate | 40 | Sigma-Aldrich | S5636 | 250x (-20°C) |
| alpha-Ketoglutarate | 5 | Sigma-Aldrich | 75892 | 250x (-20°C) |
| Citrate | 130 | Sigma-Aldrich | 251275 | 250x (-20°C) |
| Lactate | 1600 | Sigma-Aldrich | L7022 | 250x (-20°C) |
| Malate | 5 | Sigma-Aldrich | M7397 | 250x (-20°C) |
| Malonate | 10 | Sigma-Aldrich | M1296 | 250x (-20°C) |
| Pyruvate | 50 | Sigma-Aldrich | P2256 | 250x (-20°C) |
| Succinate | 20 | Sigma-Aldrich | S3674 | 250x (-20°C) |
| Creatine | 40 | Sigma-Aldrich | C0780 | 500x (-20°C) |
| Creatinine | 75 | Sigma-Aldrich | C4255 | 500x (-20°C) |
| Glutathione | 25 | Sigma-Aldrich | G6013 | 500x (-20°C) |
| Fructose | 40 | Sigma-Aldrich | F3510 | 500x (-20°C) |
| Galactose | 60 | Sigma-Aldrich | G5388 | 500x (-20°C) |

| | | | | |
|---|---|---|---|---|
| Acetone | 60 | Sigma-Aldrich | AX0120 | 5000x (-20°C) |
| Formate | 50 | Sigma-Aldrich | 94318 | 5000x (-20°C) |
| Glycerol | 120 | Sigma-Aldrich | G2025 | 5000x (-20°C) |
| Hypoxanthine | 10 | Sigma-Aldrich | H9377 | 1000x (-20°C) prepared in 0.2M HCl |
| Uridine | 3 | Sigma-Aldrich | U3003 | 1000x (-20°C) prepared in 0.2M HCl |
| Uric acid | 350 | Sigma-Aldrich | U2625 | 250x Prepared fresh in 1M NaOH |
| Urea | 5000 | Sigma-Aldrich | U5378 | 250x Prepared fresh |
| Glucose | 5000 | Thermo Fisher | 15023-021 | 100x Prepared fresh |
| ***Vitamins*** | | | | |
| Indicated vitamin concentrations are according to supplementation with RPMI 1640 100X vitamin mix (Sigma-Aldrich-Aldrich R7256) | | | | |
| Concentrations significantly different versus expected values based on metabolite profiling, as described before[3] | | | | |
| Biotin | 0.82 | | | |
| Choline | 21.49 | | | |
| Folic acid | 2.27 | | | |
| Inositol | 194.27 | | | |
| Niacinamide | 8.19 | | | |
| p-Aminobenzoate | 7.29 | | | |
| Pantothenate | 1.05 | | | |
| Pyridoxine | 4.86 | | | |
| Riboflavin | 0.53 | | | |
| Thiamine | 2.96 | | | |
| Vitamin B-12 | 0.004 | | | |
| ***Other*** | | | | |
| Phenol red | 14 | Sigma-Aldrich | P5530 | 100x (-20°C) |

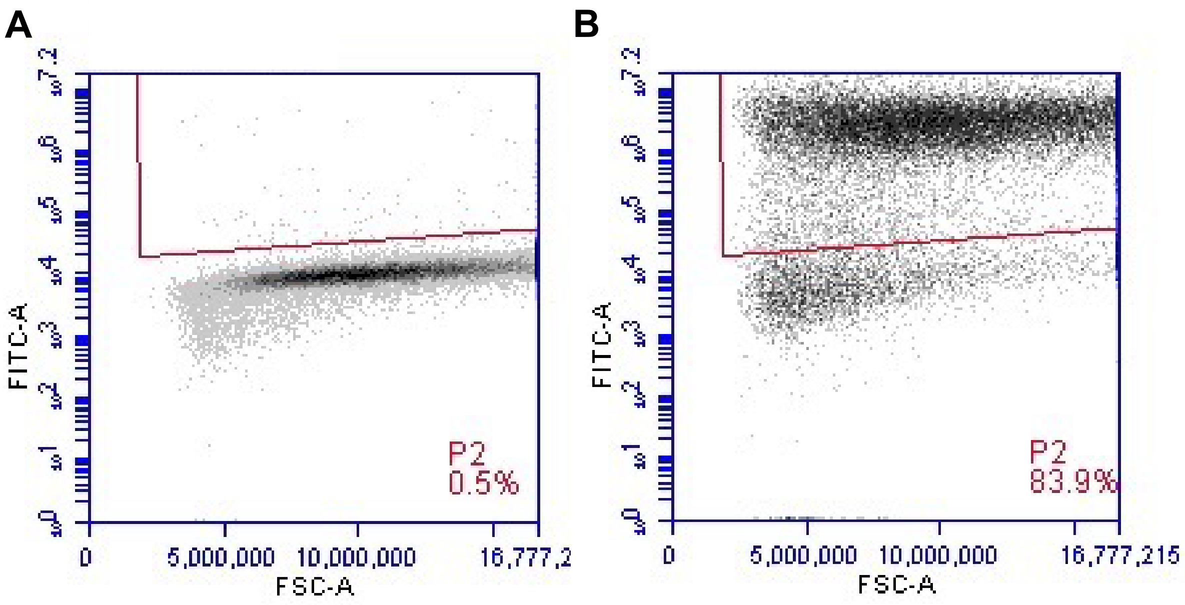


**Figure S1. Flow cytometry assessment of cardiomyocyte purity in Day 16 hPSC-CMs.** (A-B) Representative flow cytometry dot plots comparing (A) undifferentiated hPSCs (negative control) and (B) Day 16 hPSC-CMs stained for the cardiomyocyte-specific marker cTnT.

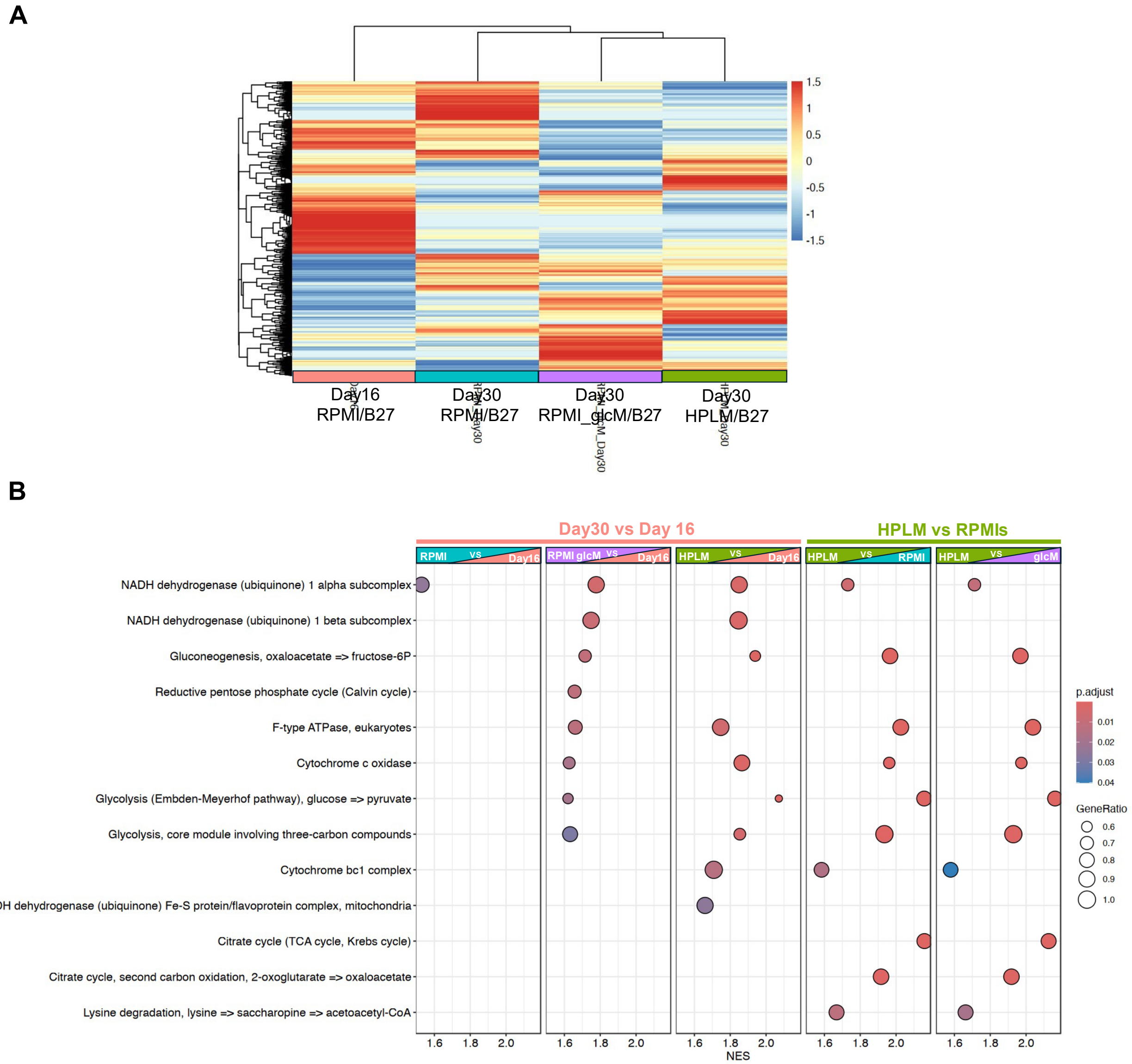


**Figure S2. Transcriptomic profiling of hPSC-CMs cultured in different media.** (A) Heatmap clustering of the top 1,000 genes with highest variance in Day 16 hPSC-CMs cultured in RPMI/B7, and Day 30 hPSC-CMs cultured in standard RPMI/B27, RPMI_glcM/B27, and HPLM/B27. (B) Dot plots of top upregulated metabolic KEGG pathways identified by Gene Set Enrichment Analysis (GSEA) across five experimental comparisons: (1) Day 30 RPMI/B27 vs. Day 16, (2) Day 30 RPMI/B27_glcM vs. Day 16, (3) Day 30 HPLM/B27 vs. Day 16, (4) Day 30 HPLM/B27 vs. Day 30 RPMI/B27, and (5) Day 30 HPLM/B27 vs. Day 30 RPMI/B27_glcM. Dot size reflects the number of genes associated with a pathway; color indicates adjusted p-value.

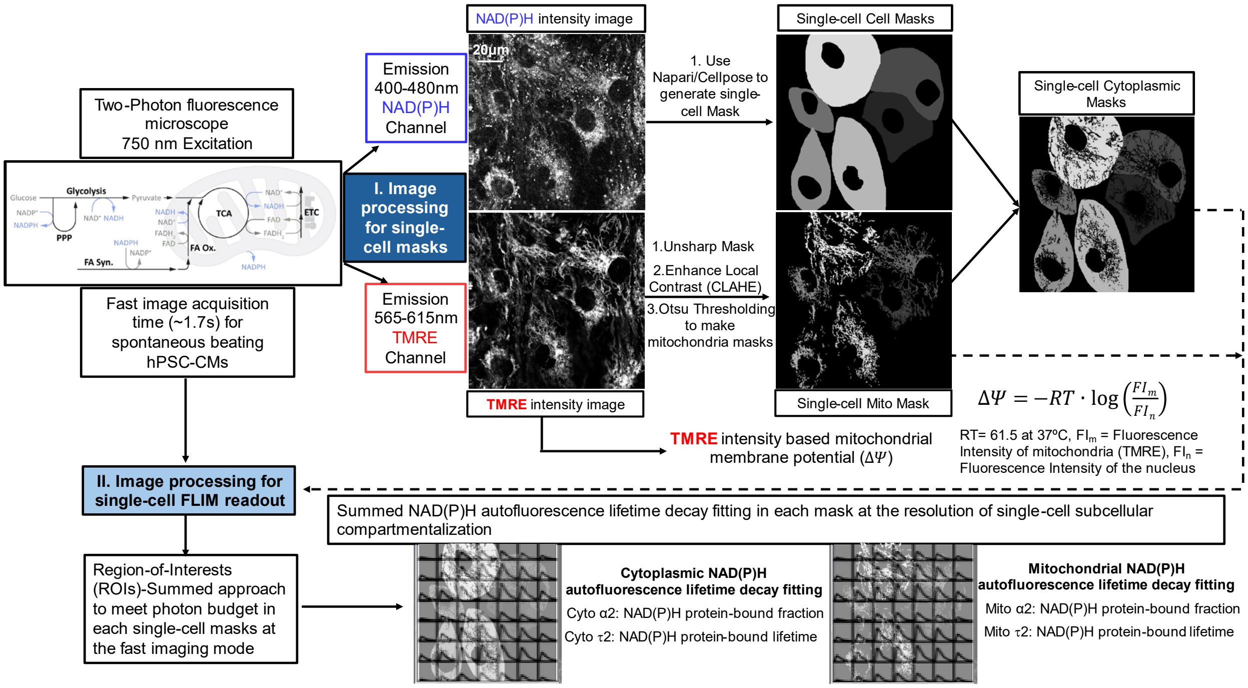


**Figure S3. Workflow of FLIM-based NAD(P)H metabolic analysis in hPSC-CMs.** (A) Single-cell segmentation and mitochondrial membrane potential (ΔΨ) quantification. Image processing workflow for generating subcellular masks (cell, mitochondria, cytoplasm) and measuring single-cell mitochondrial membrane potential (ΔΨ) by TMRE (tetramethylrhodamine, ethyl ester). TIF-formatted NAD(P)H intensity images were imported into Napari[4] and Cellpose[5] to generate single-cell whole-cell and nuclear masks. Mitochondrial segmentation was performed in ImageJ from TMRE intensity images using a custom pipeline adapted from established morphological analysis methods[6]. Mitochondrial membrane potential was quantified based on TMRE intensity in mitochondrial and normalized by nuclear intensity by equation $\Delta\Psi = -RT \cdot \log\left(\frac{FIm}{FIn}\right)$ followed by previous protocol [7]. (B) NAD(P)H fluorescence lifetime analysis. Region-of-interest (ROI)-summed approach for quantifying NAD(P)H fluorescence lifetime parameters (τ2, α2) in cytoplasmic and mitochondrial compartments. To ensure sufficient photon counts for reliable decay fitting, the ROI-Summing method was implemented as detailed in prior methodology[8]. NAD(P)H fluorescence lifetime data were processed using SPCImage software, with two-component exponential decay models ( $\mathrm{I(t)} = \alpha_1 e^{-t/t_1} + \alpha_2 e^{-t/t_2} + \mathrm{C}$ ) applied as described previously [9]. Single-cell NAD(P)H lifetime parameters (mitochondrial and cytoplasmic compartments) were extracted from SPCImage-generated ASC files using Python-based parsing and compiled into structured datasets via Excel.

### Supporting Video Legends

**Video S1**
Representative video of the contractility of Day 30 hPSC-CMs cultured from days 16 to 30 in RPMI/B27 containing 5 mM glucose.

**Video S2**
Representative video of the contractility of the Day 30 hPSC-CMs cultured from days 16 to 30 in HPLM/B27.

**Video S3**
Representative video of calcium transient of the Day 30 hPSC-CMs cultured from days 16 to 30 in RPMI/B27 containing 5 mM glucose.

**Video S4**
Representative video of calcium transient of the Day 30 hPSC-CMs cultured from days 16 to 30 in HPLM/B27.
